# *Drosophila* neural stem cells show a unique dynamic pattern of gene expression that is influenced by environmental factors

**DOI:** 10.1101/2020.12.04.411991

**Authors:** Alix Goupil, Carole Pennetier, Anthony Simon, Patricia Skorski, Allison Bardin, Renata Basto

## Abstract

With the aim of developing a sensor for chromosome loss *in vivo*, we used the well-established GAL4/GAL80 system combined with a visual GFP marker in *Drosophila*. We show a low frequency of green cells in most *Drosophila* tissues, suggesting low aneuploidy levels. Unexpectedly, in the brain, GFP positive cells are more frequent, but in this case, they do not represent chromosome loss. Using genetic manipulations, RNA FISH and time-lapse microscopy, we uncovered a dynamic and reversible silencing of *GAL80* that occurs in *Drosophila* neural stem cells. Further, we showed that this novel gene expression regulation is influenced by environmental changes such as temperature variations or food composition. These results have important implications for the *Drosophila* community, namely the possible interpretation of false positive cells in clonal experiments. Additionally, they also highlight a level of mosaicism and plasticity in the brain, consistent with possible epigenetic regulation of fly chromosomes, which is different from other organs and tissues.

## Introduction

All multicellular organisms originate from one unique totipotent cell, the zygote. This single cell divides to generate a high number of different cell types in different organs. Frequently, different cell lineages can adopt different morphologies which might be related with their function (Prasad and Alizadeh, 2019). The balance between proliferation and differentiation has to be finely tune in order to allow, during development, accurate gene expression patterns, which ensure cell fate determination (Rué and Arias, 2015). The establishment of appropriate developmental programs are tightly controlled in space and in time, such as in the optic lobe of the *Drosophila* larval brain where the combination of temporal and spatial axes in a set of neural stem cells generates highly complex neuronal diversity (Isshiki et al., 2001; Li et al., 2013; Suzuki et al., 2013; Erclik et al., 2017). Patterns of gene expression can be established by epigenetic modifications in proliferating cells, which can then be inherited by daughter cells and stably maintained over time. Genome plasticity such as the control of gene expression in response to environmental changes also influences tissue and organ behavior and development with important consequences in organism fitness (Tian and Marsit, 2018).

Most cells of a given organism present the same genetic information, which is transmitted throughout generations to maintain genetic stability. Defects in chromosome segregation result in deviations to the normal diploid chromosome content (Siegel and Amon, 2012; Bakhoum et al., 2014) In this case, cells are aneuploid, which can impact several processes such as proliferation capacity, protein homeostasis, chromosome and genetic instability (Pfau et al., 2016). Interestingly, aneuploid cells might also have different fates. In *Drosophila*, for example aneuploid wing disc cells are eliminated by apoptosis (Dekanty et al., 2012), while neural stem cells in the brain seem to lose proliferative capacity and undergo premature differentiation (Gogendeau et al., 2015). In humans, whole organism aneuploidies are frequently inviable. Developmental diseases such Down and Turner syndromes are associated with mental retardation (Lippe, 1991; Antonarakis et al., 2004). In cancer, complex aneuploidies have been described which are thought to contribute to tumour initiation and evolution (Harris and Boveri, 2008; Sansregret and Swanton, 2017; Bakhoum and Landau, 2017).

While multiple studies have addressed the mechanisms by which aneuploid cells are generated, we still lack knowledge concerning their genesis in wild type (WT) organisms. Additionally, the factors that contribute to the maintenance of aneuploid cells in tissues or that influence their fate and outcome remain to be explored. To overcome this caveat, we have established an *in vivo* method to analyse chromosome loss in a multicellular organism. This system is based on the bipartite GAL4/UAS for detection of green nuclear fluorescent protein (GFP) combined with the repressor GAL80 inserted at specific chromosome sites in *Drosophila melanogaster*. Analysis of 20 different GAL80 fly lines corresponding to different chromosome locations across the X, II and III chromosomes revealed only the presence of a low number of GFP positive cells, consistent with low levels of aneuploidy in WT proliferative epithelial tissues such as the wing disc. Strikingly however, in developing brains a large number of green cells was noticed. Using a variety of methods, including FISH and live imaging analysis, we show that, unexpectedly these cells are not aneuploid. Instead, they seem to result from a dynamic plastic gene regulation specific of the *Drosophila* brain that we further show it is influenced by environmental outputs such as food intake and temperature variations.

## Results

### A novel strategy to monitor chromosome loss based on the GAL4/GAL80 system

With the aim of developing a chromosome loss sensor in *Drosophila melanogaster* we designed a genetic tool based the principle of the loss of heterozygosity using the well-established GAL4/UAS/GAL80 system (Suster et al. 2004; Siudeja et al. 2015; Szabad, Bellen, and Venken 2012) and the expression of a fluorescent tag-green fluorescent protein (GFP fused to a nuclear localization signal (GFP-NLS). The *GFP-NLS* expression is driven by the Upstream Activating Sequence (UAS), which is a regulatory sequence activated by the transcriptional factor GAL4 (Brand and Perrimon, 1993). In the presence of the repressor GAL80, the GAL4 transcriptional activity is inhibited (Lee and Luo, 1999; Suster et al., 2004) and thus represses *GFP-NLS* expression. In this case all nuclei should appear black since they are GFP negative (GFP−). However, upon the loss of the chromosome carrying the *GAL80* sequence, *GFP-NLS* expression should be noticed through the appearance of green fluorescence, GFP positive (GFP^+^) nuclei (Figure 1A). We wanted to develop this system to monitor random chromosome loss of any of the major autosomes (chromosomes II and III) and the X chromosome.

**Figure 1:**
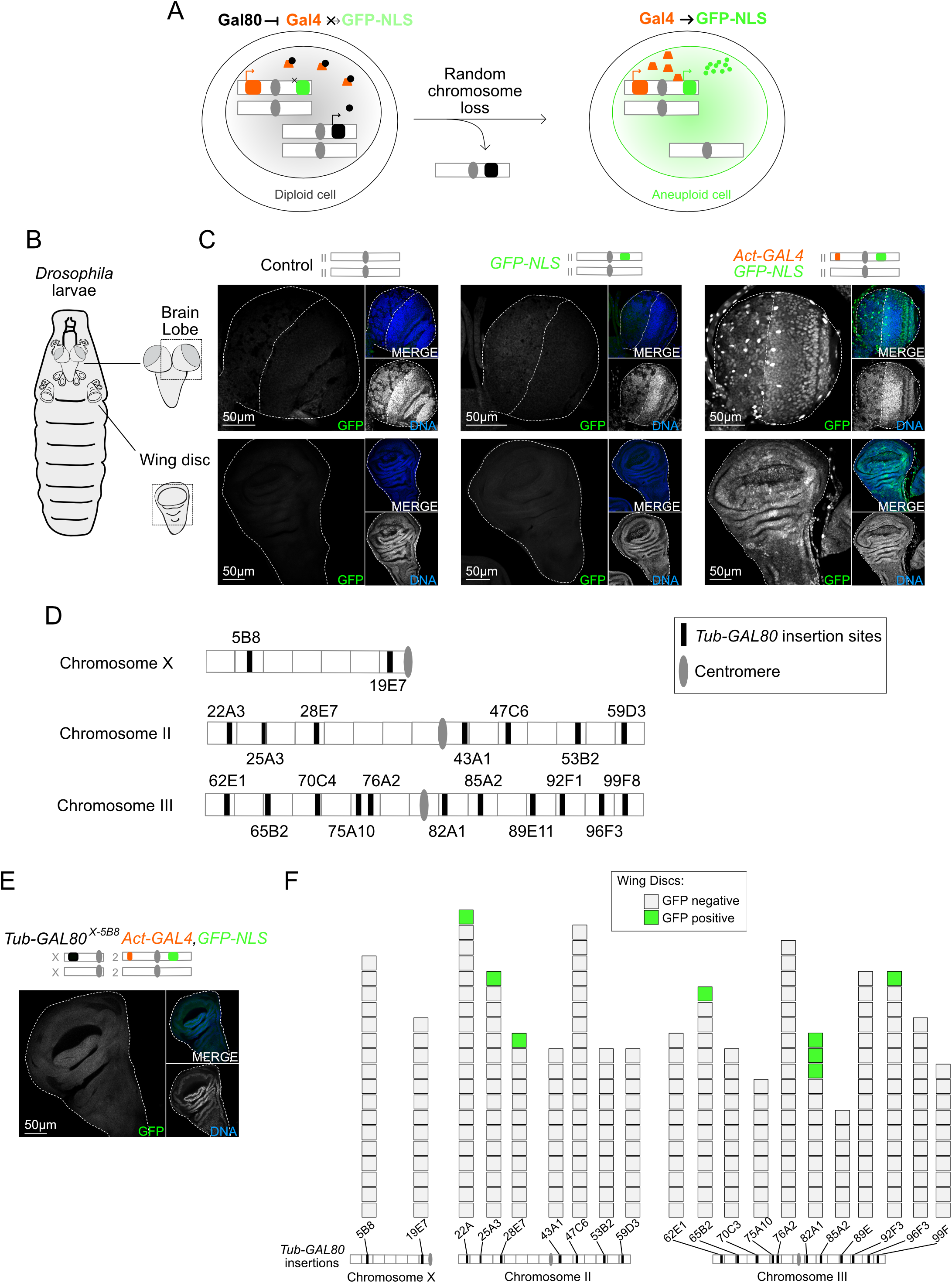
A novel strategy to monitor chromosome loss based on the GAL4/GAL80 system which is functional in *Drosophila* larval wing discs. (A) Schematic representation of the genetic system to monitor chromosome loss in *Drosophila* cells. On the left-the presence of Gal80 inhibits Gal4 and so the nuclei are black. On the right-Upon random loss of the *GAL80* containing chromosome, Gal4 is released from Gal80 repression and promotes *GFP-NLS* expression. Thus, aneuploid cells appear with green nuclei. (B) Schematic representation of the brain and imaginal discs of the *Drosophila* L3 larvae. (C) Images of whole mount brain lobes and wing discs labeled with GFP booster (grey and green in large and small insets, respectively) and DAPI for DNA (grey and blue in small insets). Control and GFP-NLS tissues present no GFP signal. In the presence of the activator Gal4, *GFP-NLS* is expressed and all cells present GFP positive (GFP^+^) nuclei. Schematic representation of the genotypes is shown above the images: *GFP-NLS* and *Act-GAL4* sequences are represented with green and orange rectangles on white chromosomes, respectively. White dotted lines delimitate tissues. (D) Representative map of the *Tub-GAL80* insertion sites on *Drosophila* chromosomes X, II and III used in this study. (E) Images of whole mount wing disc labeled with GFP booster (grey and green in large and small insets, respectively) and DAPI for DNA (grey and blue in small insets). In the presence of Gal80, Gal4 is repressed and all cells are GFP negative (GFP^−^). Schematic representation of the genotype is shown above the images: *GFP-NLS, Act-GAL4* and *Tub-GAL80* sequences are represented with green, orange and black rectangles on white chromosomes, respectively. White dotted line delimitates the wing disc. (F) Graph summarizing the results from the screen of all *Drosophila* lines carrying one copy of the *Tub-GAL80* cassette on the chromosomes X, II and III. Insertion sites are represented in the scheme bellow. Each square of the graph represents one wing disc that presented no GFP signal (grey) or at least one GFP^+^ cell (green) (n=7 to 20 wing discs /GAL80 condition).

Because chromosome loss is known to occur after chromosome mis-segregation in mitotically active cells (Levine and Holland, 2018), we focused on two *Drosophila* tissues known to be highly proliferative at third instar larvae (L3): the brain lobes, which make part of the fly central nervous system and epithelial wing discs, which are the primordial fly wings (Figure 1B).

Using meiotic recombination in the female germline, we established an *Act-GAL4, GFP-NLS Drosophila* recombinant line carrying on chromosome II the *UAS-stinger* sequence (Stable insulated nuclear eGFP) (Barolo et al., 2000) and the *GAL4* gene controlled by the ubiquitous *Actin5C* promotor (Struhl and Basler, 1993; Ito et al., 1997). Importantly, we confirmed that *Act-GAL4, GFP-NLS* expressing larvae show GFP^+^ nuclei in both brain lobes and wing discs (Figure 1C). For the establishment of the *GAL80* lines, we designed a new vector expressing a *GAL80* version which has been optimized for *Drosophila* codon usage (Pfeiffer et al., 2010) under the control of the ubiquitous *Tubulin 1α* promotor – *Tub-GAL80* (O’Donnell et al., 1994; Lee and Luo, 1999). Even though this GAL80 optimized codon was shown to have a higher GAL4 suppression capacity (Pfeiffer et al., 2010), it is known that the chromosomal environment and so its insertion site impacts transgene expression (Elgin and Reuter, 2013). To ensure the best optimal *Tub-GAL80* expression conditions, we generated a total of 20 *Drosophila* lines. Each line carried one copy of the *Tub-GAL80* transgene inserted at one specific site and it will be referred to as *Tub-GAL80* followed by the insertion site in superscript. As an example, the *Tub-GAL80* insertion at the 5B8 location on the X chromosome will be referred as *Tub-GAL80^X-5B8^*. We obtained lines with *Tub-GAL80* insertions at different locations on chromosomes X, II and III. A list summarizing the insertion sites and their position relative to the centromere can be found in Figure 1D.

### The large majority of wing disc cells are GFP negative

As a proof of concept, we screened all *Tub-GAL80* stocks for their capacity to repress the *Act-GAL4, GFP-NLS* driver. We analysed by immunofluorescence 7 to 20 wing discs per *Tub-GAL80* insertion. For all lines, the vast majority of wing discs were GFP− (361/369 in total) (Figure 1E-F). In 5 out of the 8 wing discs that contained GFP^+^ cells, these cells were restricted to only a few cells within the whole GFP− tissue (5 wing discs from 5 different *Tub-GAL80* constructs) (Supplementary Figure 1A). Interestingly, only one single *Tub-GAL80* line-*Tub-GAL80^III-82A1^*-had a high number of GFP^+^ cells (Supplementary Figure 1B). Importantly, however, this only occurred in 3 out of 12 wing discs analysed for this line (Figure 1F). This suggests that in the wing disc, most *Tub-GAL80* expressing flies, can repress *GFP-NLS* expression. The same observation, namely the absence of GFP signal, was obtained after the analysis of other imaginal discs (Supplementary Figure 1C). Overall these results showed that for 19 out of 20 lines, the system is functional with GAL80 repressing GAL4 activity in epithelial larval discs.

### The brain presents a high number of GFP^+^ cells

The larval brain lobe can be divided in two main regions, the central brain and the optic lobe. The central brain is composed of the neural stem cells, also called neuroblasts (NBs) that divide asymmetrically to self-renew and give rise to smaller and more committed progenitors, the ganglion mother cells (GMCs). Two populations of larval NBs have been identified. Type I NBs, which divide asymmetrically to give rise to GMCs that will divide once more before undergoing differentiation to generate either glia or neurons. Type II NBs generate intermediate progenitors through asymmetric cell division, which will then generate GMCs followed by differentiation (Bello et al., 2008; Boone and Doe, 2008; Homem and Knoblich, 2012). The optic lobe comprises two proliferative centers, the inner and the outer, which correspond to a pseudo-stratified epithelium called the neuroepithelium (NE). NE cells give rise to neurons necessary for the development of the visual system of the fly. Further, perineural and sub-perineural glial cells with large nuclei are found at the superficial layer of the brain (Pereanu et al., 2005). Interestingly, all these cell types are easily distinguishable by their morphology, position within the tissue and expression markers (Figure 2A-E).

**Figure 2:**
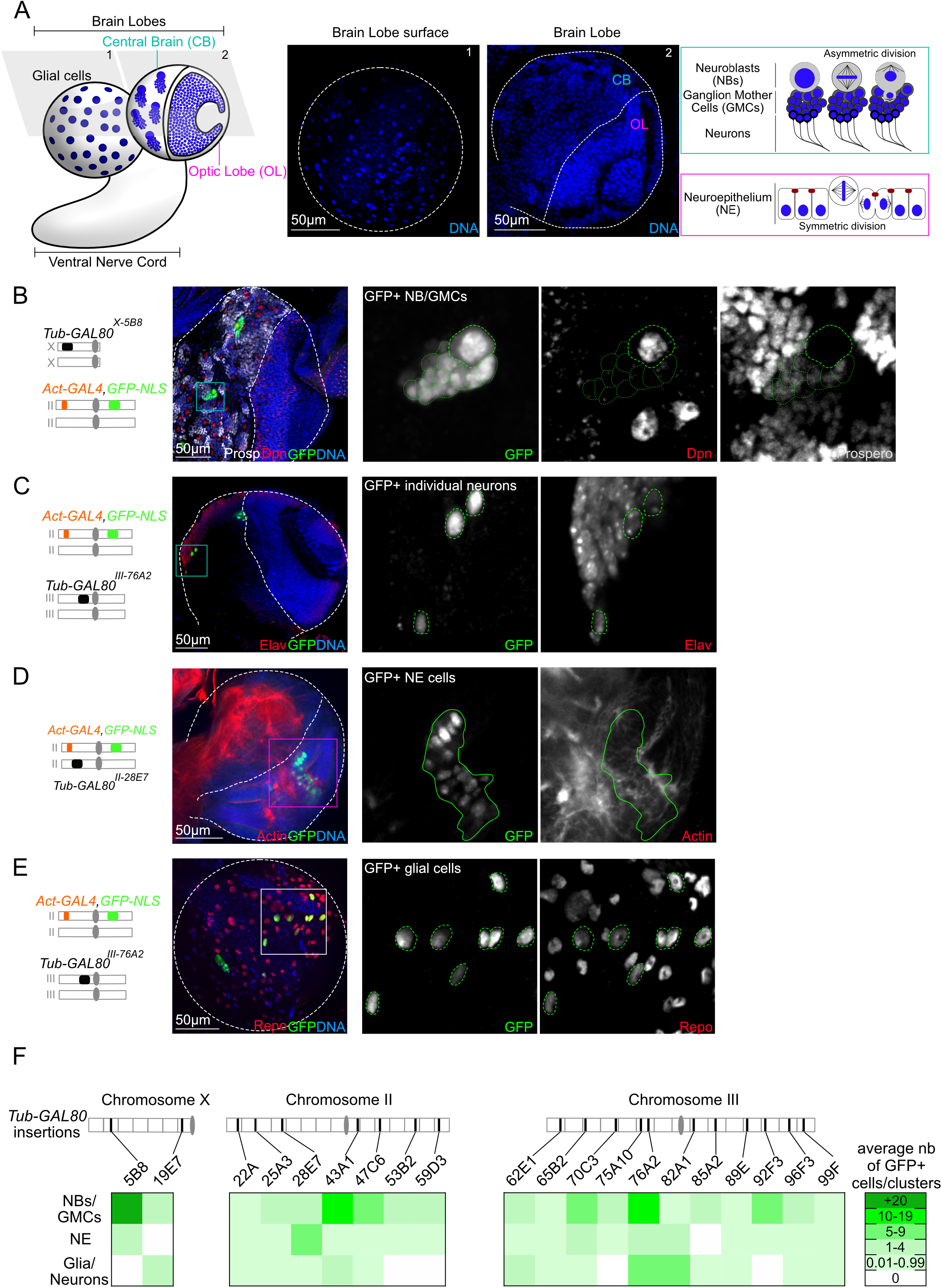
GFP positive cells correspond to different cell types in the larval brain and their frequency is variable. (A) Schematic representation of the *Drosophila* larval brain and its different cell types. Representative images of different z stacks of the brain lobes are shown with DNA in blue. Glial cells with large nuclei are present at the surface of the brain lobe (image 1). The core of the brain lobe is divided in two parts, the central brain (CB) and the optic lobe (OL) (image 2). The CB is composed of neuroblasts (NBs) that divide asymmetrically to generate ganglion mother cells (GMCs) that will then give rise to neurons. The OL is composed of neuroepithelial (NE) cells that divide symmetrically. (B-E) Images of whole mount brain lobes labeled for GFP (grey and green) and for specific markers of the different cell types (red and grey). DNA is shown in blue. Schematic representation of the genotypes is shown next to the images. Zoom insets are highlighted by colored squares. White dotted lines delimitate brain lobes and separate CB and OL. Green dotted and continuous lines surround GFP^+^ cells and clusters, respectively. GFP^+^ cells are (B) Dpn^+^ NBs with Dpn^−^/Prospero^weak^ GMCs, (C) individual Elav^+^ neurons, (D) cells of the NE which is distinguishable by the specific F-actin organization, and (E) Repo^+^ glial cells. (F) Heat map showing the average number of GFP^+^ cells/clusters (green color code) for each cell type of the brain lobe and per GAL80 condition. The absence of any green cell is represented in white, while the presence of 1 to 4 cells/clusters, 5 to 9 cells/clusters, 10 to 19 cells/clusters and more than 20 cells/clusters were represented in increased shades of green. The map of *Tub-GAL80* insertion sites is schematized above the graph.

As described above, we screened the *Tub-GAL80* lines for their capacity to repress GAL4 and thus the expression of *GFP-NLS*. Interestingly and in contrast with the results obtained for wing discs, brain lobes frequently contained GFP^+^ cells. Moreover, the morphology of the GFP^+^ cells was different among the various *Tub-GAL80* lines, suggesting that GFP^+^ cells belong to different cell populations (Figure 2B-E). The different cell types that populate the brain lobes are morphologically distinguishable and recognised through DNA labelling (DAPI) and by their spatial position within the brain (Figure 2A). Using specific markers, we confirmed by immunofluorescence the various identities of GFP^+^ cells. Indeed, Dpn^+^ NBs in the central brain occupy the first layers of the tissue, just below the surface and have large nuclei. Dpn^-^/Propero^weak^ GMCs are small nuclei juxtaposed to NBs (Figure 2A-B and Supplementary 2A). The individual Elav^+^ neurons recognised by the small-sized nuclei are dispersed in the central brain and localised deeper in the tissue (Figure 2C and Supplementary 2B). The NE, which is recognizable by the actin organization of this pseudo-stratified epithelium, is localised in the optic lobe region and NE cells contain highly arranged and small nuclei (Figure 2D and Supplementary 2C). Finally, Repo^+^ glial cells are located at the periphery of the lobes and have large nuclei (Figure 2E and Supplementary 2D). Interestingly, some brain lobes presented a mix population of GFP^+^ cells, while in others these cells were absent (Supplementary Figure 2E-F).

To obtain a quantitative view of the population of GFP^+^ cells, we counted and categorized GFP^+^ cells into the following subtypes: 1) GFP^+^ clusters which included NBs and associated GMCs (independently of their number); 2) GFP^+^ clusters for NE cells and 3) individual GFP^+^ cells for differentiated neurons and glia. We represented with a colour code the number of GFP^+^ cells/clusters, taken as independent groups. Interestingly, analysis of 459 brain lobes from all the *Tub-GAL80* lines (minimum of 14 brain lobes per *Tub-GAL80* insertion line) revealed that the number and identity of GFP^+^ cells varied between different *Tub-GAL80* lines and even more surprisingly within the same *Tub-GAL80* line (Supplementary Figure 3). For example, *Tub-GAL80^X-5B8^* on the X chromosome presented a high number of green NBs and associated GMCs, but no green neurons or glial cells were detected and only very few green NE cells were observed. (Supplementary Figure 2A and Supplementary Figure 3A). In contrast, *Tub-GAL80^III-82A1^* on chromosome III, showed a high number of green neurons and glia and only a low number of green NBs/GMCs (Supplementary Figure 3E), while *Tub-GAL80^II-22A^* (chromosome II) displayed only a few green cells in each category (Supplementary Figure 2F and Supplementary Figure 3B).

To be able to compare the frequency and identity of green cells, we plotted the mean value of each group category (cell identity) for all the *Tub-GAL80* lines (Figure 2F). Strikingly, this analysis showed that green cells were highly frequent in the central brain of all the *Tub-GAL80* lines, with the highest incidence in the NB and GMC cell population. It is important to mention that we did not find a trend in the position or spatial arrangement of GFP^+^ NBs or even other cell types within the different brain lobes analysed. These observations suggest that there is no particular stereotype or spatial pattern of the cells expressing *GFP-NLS*, but rather that they are randomly positioned.

To use an alternative means of quantifying the frequency of GFP^+^ cells in the brain, we measured the area of the GFP signal and express it as the percentage of coverage per brain lobe area (Supplementary Figure 4A). As expected, the high frequency revealed by the colour code correlated with highest coverage values (Figure 2F and Supplementary Figure 4B). We conclude that a high number of GFP^+^ cells including NBs, GMCs, NE cells, glia and neurons are present, independently of the *Tub-GAL80* insertion site. This result highly contrasts with the findings in wing discs using the same *Tub-GAL80* lines.

### GFP positive cells in the brain are not the by-product of chromosome loss

We next focused our analysis on understanding the origin of the difference in the frequency of GFP^+^ cells between wing discs and brains. An obvious hypothesis to explain the high incidence of GFP^+^ cells in the brain was aneuploidy due to the loss of the *GAL80* containing chromosome (Figure 1A). We reasoned that if this was the case, GFP^+^ cells should be aneuploid in contrast to diploid GFP− surrounding cells. To assess the ploidy of the GFP^+^ cells, we used a Fluorescent *In Situ* Hybridization (FISH) protocol using probes against the chromosome carrying the *Tub-GAL80* insertion. As reported previously, dotty FISH signals can be easily noticed ranging from 1 to 4 structures, which correspond most likely to different degrees of chromosome pairing and unpairing (Joyce et al., 2012). While this limitation precluded the use of FISH signals for precise ploidy assessment, we did notice that the FISH signals were similar between GFP^+^ and GFP− cells (Figure 3). Thus, this argues against the GFP^+^ cells being aneuploid via chromosome loss, which would predict an overall decrease in the number of FISH signal dots. Additionally, to avoid possible mis-interpretation due to chromosome pairing, typical of *Drosophila* cells, we examined FISH signals in mitotic NBs which facilitates the analysis of FISH probes combined with individual chromosomes (Gatti et al., 1994). We confirmed that GFP^+^ cells were not aneuploid as they presented the same number of FISH dots (Figure 3B) than control GFP− NBs. These results suggest that the high frequency of green cells observed across different *Tub-GAL80* lines do not result from chromosome loss.

**Figure 3:**
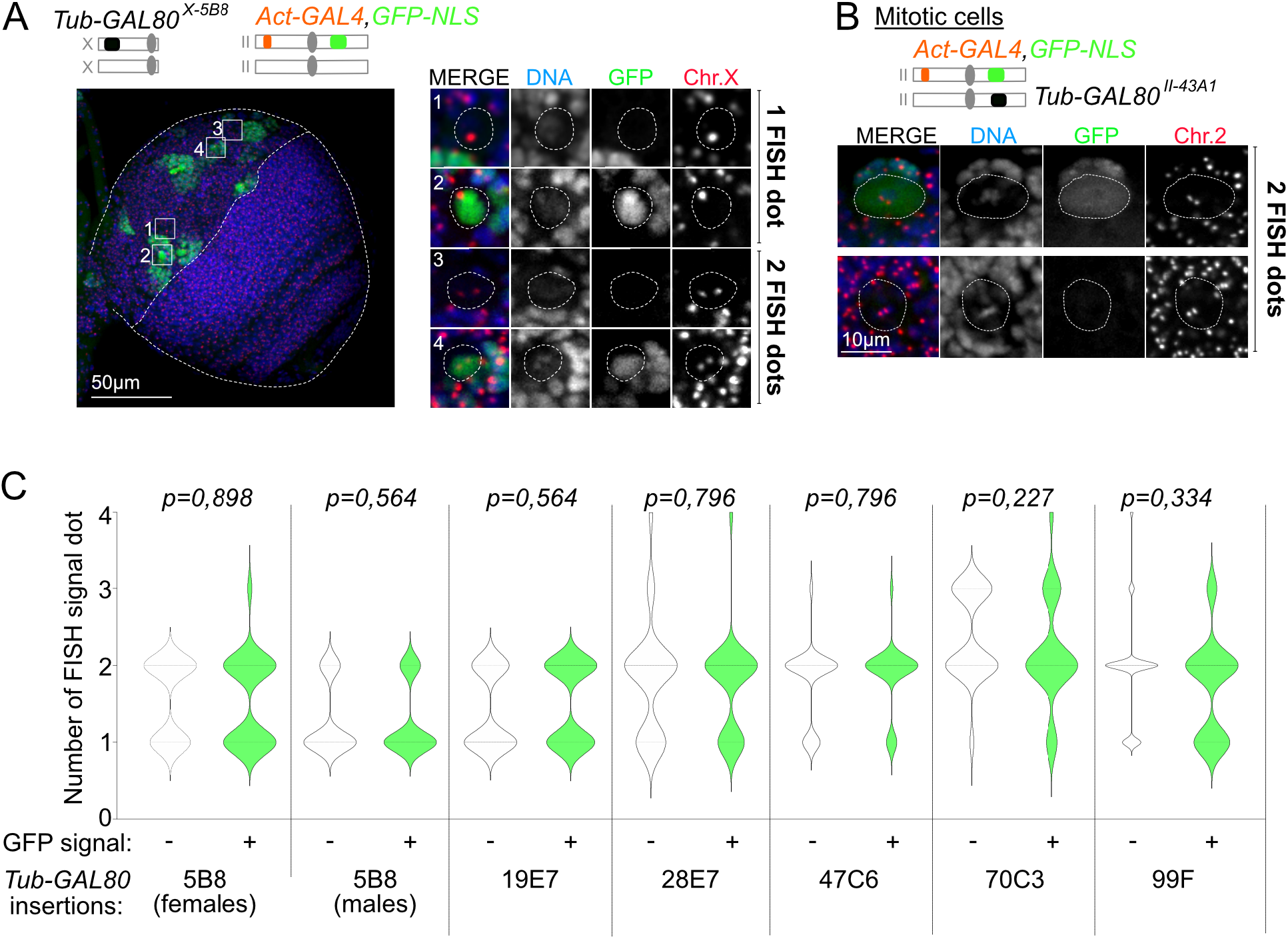
GFP positive cells are not aneuploid in the larval brain. (A-B) Fluorescent *in situ* hybridization with probes for the (A) X chromosome or (B) chromosome II (red and grey in zoom insets) combined with labeling with GFP booster (green and grey in zoom insets) and DAPI for DNA (blue and grey in zoom insets) of the brain lobe. The number of FISH signal dot is similar between GFP^+^ and GFP^−^ (A) interphase and (B) mitotic NBs. Schematic representation of the genotypes is shown above the images. White dotted lines surround brain lobe and NBs in large and zoom insets, respectively. (C) Violin plot representing the number of FISH signal dots between GFP^+^ and GFP^−^ cells. FISH signals correspond to the chromosomes X, II or III for conditions where *Tub-GAL80* was inserted at positions 5B8 (n=43 cells for females and n=50 cells for males) and 19E7 (n=47 cells), 28E7 (n=50 cells) and 47C6 (n=50 cells) or 70C3 (n=47 cells) and 99F (n=60 cells), respectively. FISH signals are variable between conditions but similar between GFP^+^ (green) and GFP^−^ (white) cells from the same condition. Statistical non-significance was determined by a Mann-Whitney test and p corresponds to the p-value.

### Mitotic recombination does account for the presence of GFP positive cells in the brain

Non-sister chromatids can exchange chromosome pieces by cross-over in somatic cells, an event called mitotic recombination. Such event has been previously described in *Drosophila* (Stern, 1936; Kaplan, 1953; Siudeja et al., 2015; Siudeja and Bardin, 2017). We hypothesised that after replication, mitotic recombination might explain the loss of *Tub-GAL80* heterozygosity. In light of this scenario after cell division one daughter cell should be homozygous for *Tub-GAL80* (GFP−) and its sister cell should lack the *Tub-GAL80* sequence (GFP^+^) (Supplementary Figure 5A). We first tested this hypothesis considering the X inserted GAL80 line – *Tub-GAL80^X-5B8^*. We compared GFP-NLS signals between males and females as mitotic recombination cannot occur between non-homologous X and Y chromosomes. Interestingly, both male and female larval brains presented similar levels of GFP^+^ cells (Supplementary Figure 2A and Supplementary Figure 5B and H), suggesting that at least for the X chromosome mitotic recombination does not account for loss of the repressor GAL80. We next tested other genetic conditions where mitotic recombination was inhibited by the use of balancer chromosomes. Balancer chromosomes contain chromosomal inversions that supress crossing over with the homologous chromosome (Novitski and Braver, 1954). We used FM7, CyO and TM6 fly lines, which are specific balancers for the X, II and III chromosomes, respectively. Interestingly, the frequency of brain lobes presenting GFP^+^ cells was unchanged using any of the balancer chromosomes when compared to controls (Supplementary Figure 5C-H). Additionally, we did not observe an obvious decrease in the GFP coverage when mitotic recombination was inhibited, as values were highly variable between and within conditions as in control experiments (Supplementary Figure 5I and Supplementary Figure 3). Altogether, this suggests that the presence of GFP^+^ cells and thus, the expression of *GFP-NLS* specifically in the brain does not result from mitotic recombination.

### Analysis of frequently used *Tub-GAL80* lines confirms the lack of GAL80 inhibition in *Drosophila* NBs and alert on its use for MARCM analysis

Our results so far suggest that the high incidence of green cells found in *Drosophila* brains cannot be explained by chromosome loss or mitotic recombination. Thus, we hypothesised that in GFP^+^ cells, the *Tub-GAL80* sequence was still present but likely not expressed. One trivial explanation for our results was that the *Tub-GAL80* sequences generated and used in this study may contain a particular feature that precludes its use in the brain. To test this possibility, we analysed *GAL80 Drosophila* lines with different origins and established by other labs available from the Bloomington *Drosophila* Stock Center. We first tested three different lines containing one copy of *Tub-GAL80* inserted on the X chromosome (*#5132-Tub-GAL80^X-1C2^*) and chromosome III (*#5191-Tub-GAL80^III-75E1^* and *#9490-Tub-GAL80^III^*). Interestingly, similar to our *Tub-GAL80* lines, the large majority of brain lobes presented high levels of GFP^+^ cells in contrast to the absence or low levels of GFP-NLS signals in the wing discs (Figure 4A-D and G). These results suggest that like our *Tub-GAL80* lines described above, the *Tub-GAL80* lines available from other labs cannot inhibit GAL4 activity in the brain, arguing against a specific effect of our lines.

**Figure 4:**
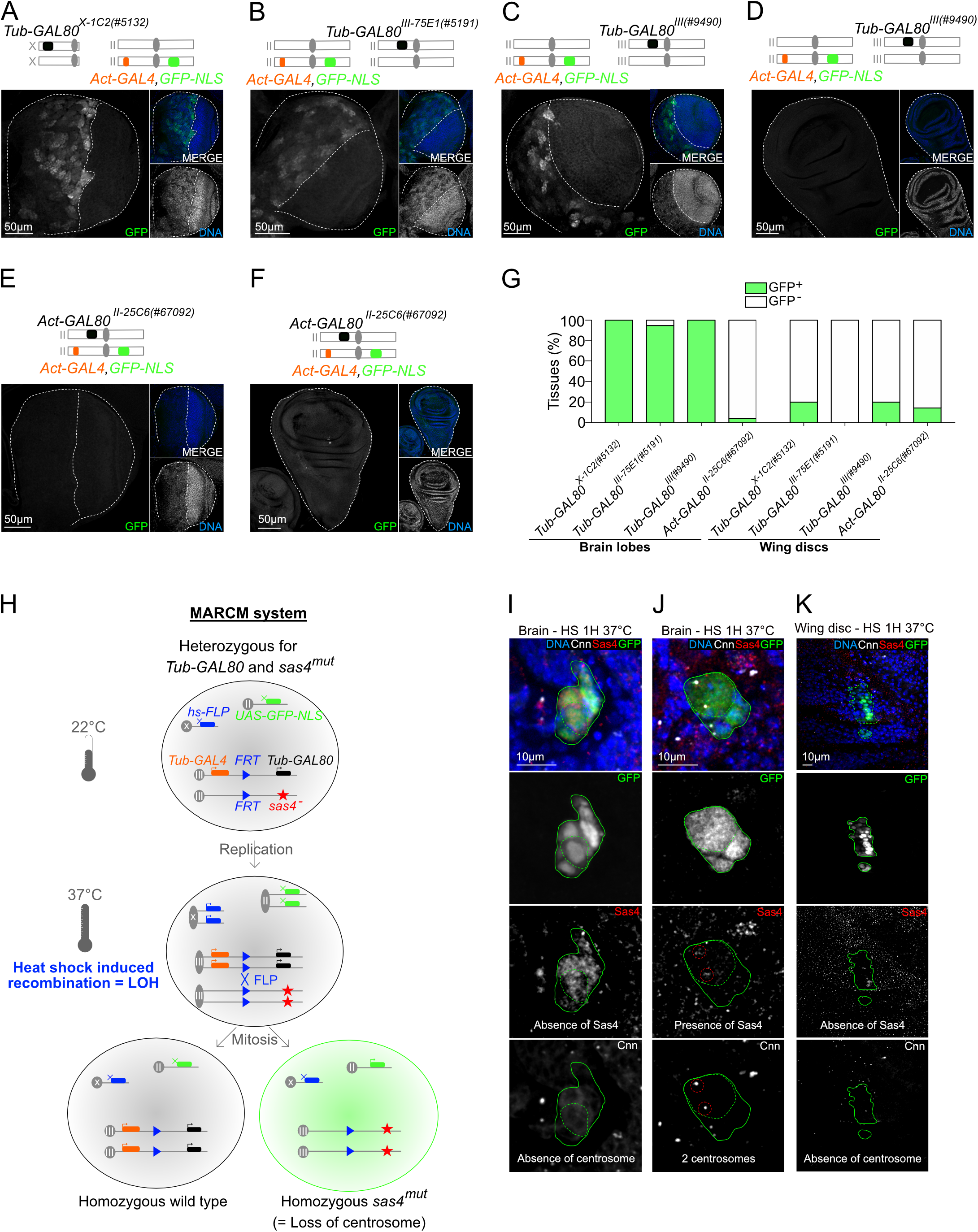
The *Tub-GAL80* expression is not sufficient to induce a complete GAL4 repression, precluding its use in MARCM analysis. (A-F) Images of whole mount (A-C and E) brains lobes or (D and F) wing discs labeled with GFP booster (grey and green in large and small insets, respectively) and DAPI for DNA (grey and blue in small insets). White dotted lines delimitate tissues. Schematic representation of the genotypes is shown above the images. (G) Graph bar showing the percentage of tissues without (white) and with GFP signal (green). (H) Schematic representation of the MARCM system. In principle, after recombination of FRT sites by the heat-induced FLP recombinase, the daughter cells lose their heterozygosity. One cell becomes homozygous mutant and labeled with GFP due to the concomitant loss of the *Tub-GAL80* sequence. The other cell becomes homozygous wild type and it is unlabeled. (I-K) Images of GFP^+^ clones in (I-J) brain lobes and (K) wing disc of *hs-FLP/+, UAS-GFP-NLS/+; Tub-GAL4,FRT82B,Tub-GAL80/FRT82B,sas4^mut^* flies heat-shocked at 37ºC for 1 hour. Green dotted and continuous lines surround GFP^+^ NBs and clones, respectively. (I) *Sas4* mutant GFP^+^ NB without centrosome. (K) Wild type GFP^+^ NB with two centrosomes. (K) *Sas4* mutant GFP^+^ clones in the WD.

In the experiments described so far, *GAL4* and *GAL80* expression are under the control of different promoters, even if both are strong and ubiquitous - actin and tubulin, respectively. We decided to test if differences in the promoters could contribute to inefficient repression of GAL4 as previously reported (Pfeiffer et al., 2010). Strikingly, this was indeed the case as very few GFP^+^ cells were noticed in the *Drosophila* line carrying the *GAL80* sequence under the control of an *Actin* promotor - *Act-GAL80* (#67092) (Figure 4E-G). It is important to notice that in contrast to the other *Tub-GAL80* lines inserted on chromosomes X and III and used in this study, the *Act-GAL80* sequence is located at position 25C6 on chromosome II. This position might also impact *GAL80* expression levels and explain the efficiency in GAL4 inhibition and thus, *GFP-NLS* repression. This suggests that both the chromosomal insertion and the use of the same promoters in repression/activation experiments accounts for stoichiometry between *GAL4* and *GAL80* expression levels.

The GAL4/GAL80 system is widely and routinely used by the *Drosophila* community to control gene expression. Importantly, this system has been used in MARCM experiments for neuronal lineage tracing in the developing *Drosophila* brain (Lee and Luo, 1999; Ren et al., 2016). This is based on the loss of heterozygosity (LOH) after mitotic recombination by the heat-induced FLP recombinase at specific FRT sites. LOH generates labelled mutant clones that lack the *GAL80* sequence and unlabelled wild type cells homozygous for *GAL80* (Figure 4H). We wanted to ensure that GFP^+^ clones generated in the MARCM experiment was only due to LOH and not *GAL80* repression as we described above in the larval brain. We used *heat-shock FLP;; FRT^82B^sas4^mut^* expressing flies crossed with *UAS-GFP-NLS; Tub-GAL4, FRT^82B^, Tub-GAL80*. The *sas*-4 gene encodes for a protein essential for centriole duplication and *sas4* mutant cells lack centrosomes (Basto et al., 2006). Heat-shock at 37°C for 1H was used to induce FLP mediated recombination. We analysed GFP^+^ clones in L3 brains and wing discs. As expected, we observed GFP^+^ mutant clones without centrosomes (Figure 4I). Surprisingly, we also observed GFP^+^ clones containing centriolar staining of Sas-4 protein, indicating that the WT sas4 gene was present (Figure 4J). Most likely, these latter clones were not generated through *GAL80* sequence loss upon FLP/FRT-mediated LOH, but they rather represent GFP^+^ cells arising in the manner described above. Interestingly, and as a control, we also analysed brains that were not heat-shocked and again observed GFP^+^ clones, even if at lower frequencies (Supplementary Figure 6A-B). Importantly, however in the wing discs in contrast to the brains, we only observed GFP^+^ clones after heat-shock induction and these clones were *sas-4* negative (Figure 4K and Supplementary Figure 6C).

Importantly in this set of experiments, we found false positive cells using the MARCM system in the larval brain as it presents a mix population of GFP^+^ clones generated both by the loss of *GAL80* sequence or by the loss of *GAL80* expression. To our knowledge, a brain-specific *GAL80* expression regulation has never been described before.

### Increasing *Tub-GAL80* levels at certain chromosome locations efficiently inhibits GAL4 activity in the brain

It has been shown that one limitation of the GAL4/GAL80 system results from the stoichiometry in the system, since if both activator and repressor are expressed at the same levels, one copy of *GAL80* might not be sufficient to repress GAL4 (Pfeiffer et al., 2010). In our experimental set up, *Act-GAL4* and *Tub-GAL80* are both ubiquitously expressed with strong actin and tubulin promoters, respectively. In principle, one copy of *Tub-GAL80* should be sufficient, especially because it has been optimized for *Drosophila* codon usage. However, the high frequency of GFP^+^ cells observed in brain lobes and described above raise the question whether increased levels of GAL80 would favour GAL4 inhibition in the brain.

We built 5 different *Drosophila* recombinant lines harbouring two copies of *Tub-GAL80* inserted at two different and distant loci of the same chromosome and will be referred to as *2xTub-GAL80* (Supplementary Figure 7A). Interestingly, all brain lobes (n=50) from two lines containing *2xTub-GAL80* insertions on chromosomes II (*2xTub-GAL80^II-22A,53B2^*) and III (*2xTub-GAL80x2^III-62E1,96F3^*) were GFP− (Figure 5A and D-E). Surprisingly, however, the two other lines for the same chromosomes (*2xTub-GAL80^II-25A3,59D3^* and *2xTub-GAL80^III-65B2,99F^*) presented GFP^+^ cells, though with a highly reduced frequency (Figure 5D-E and Supplementary Figure 7B-C). These results suggested that the addition of one extra-copy of *Tub-GAL80* on chromosomes II and III increased the capacity of GAL80 to inhibit GAL4 activity. Moreover, they also show that the position of the *Tub-GAL80* insertion influences its capacity to supress *GFP-NLS* expression, or in other words that *GAL80* expression might be conditioned by its position within the genome.

**Figure 5:**
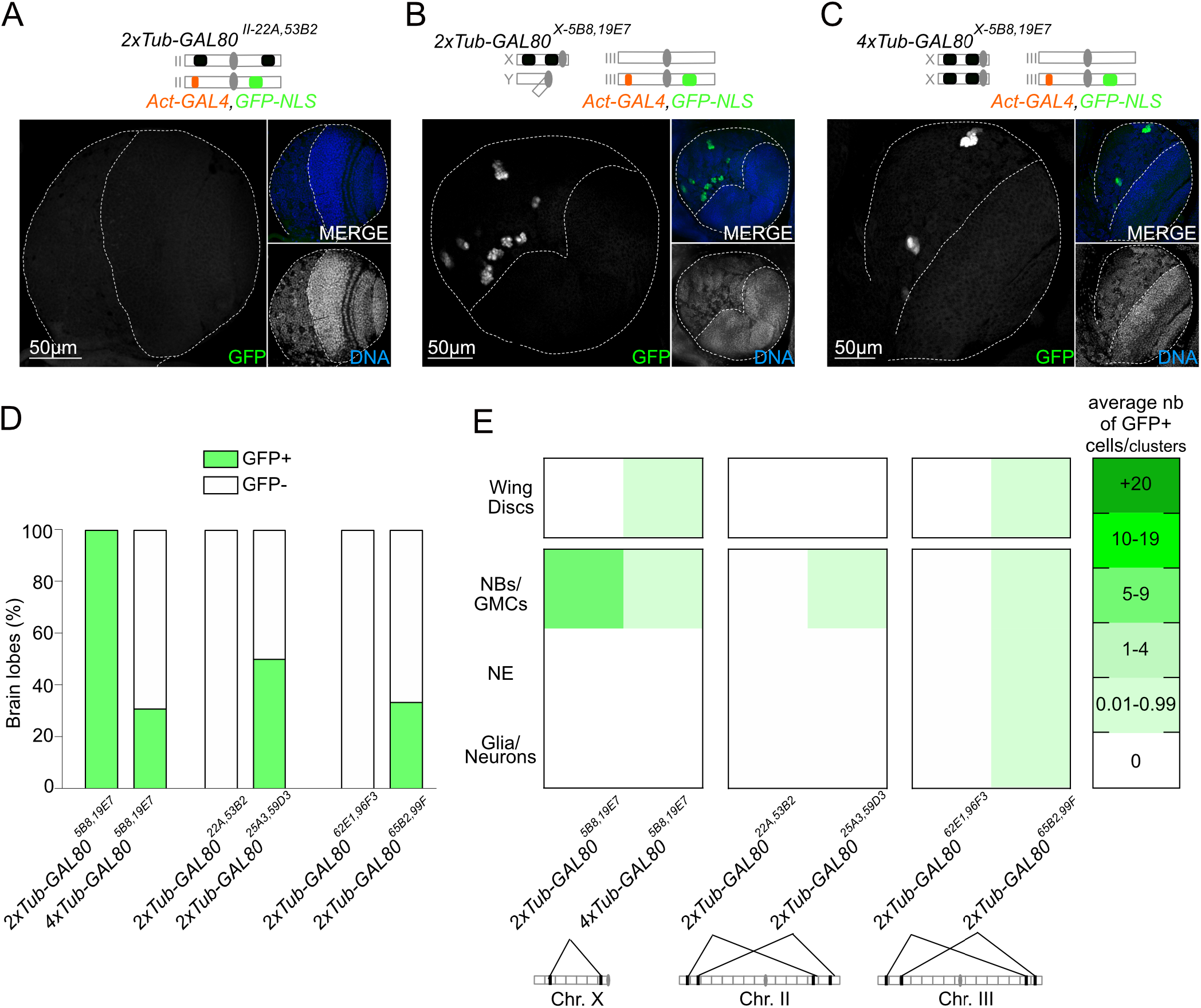
The *GAL80* levels partially explain GFP signal appearance. (A-C) Images of whole mount brain lobes labeled with GFP booster (grey and green in large and small insets, respectively) and DAPI for DNA (grey and blue in small insets). White dotted lines delimitate brain lobes. Schematic representation of the genotypes is shown above the images. (D) Graph bar showing the percentage of brain lobes without (white) and with GFP signal (green) (n=22 to 26 brain lobes/condition). (E) Heat map summarizing the average number of GFP^+^ cells/clusters per GAL80 condition in wing discs (n=7 to 18 wing discs/condition) and for each cell type in brain lobes (n=22 to 26 brain lobes/condition). The *Tub-GAL80* insertions map is schematized above the graph.

We next analysed brains containing the recombinant *2xTub-GAL80* insertions on the X chromosome - *2xTub-GAL80^X-5B8,19E7^*, also confirmed by PCR (Supplementary Figure 7D). Strikingly, a high number of GFP^+^ NBs and GMCs was noticed in this condition, either in the presence of two copies (*2xTub-GAL80^X-5B8,19E7^* heterozygous) or even four copies (*2xTub-GAL80*^-5B8,19E7^* homozygous) (Figure 5C-E). Importantly, the vast majority of wing discs were still GFP- (Supplementary Figure 7E and Figure 5E). To test if a NB specific promoter resulted in the change of GFP^+^ NBs frequency in the *2xTub-GAL80^X-5B8,19E7^* we used the NB specific promotor *Worniu* to drive *GAL4* expression. However, even in this condition, GFP^+^ cells were very frequent (Supplementary Figure 7F).

Taken together our results show that increasing GAL80 levels results in a more effective inhibition of GAL4 in the brain for certain chromosomes and chromosome territories with the X chromosome representing an exception. Interestingly, our work revealed that *GAL80* insertions on the X chromosome behave quite differently from other lines, suggesting that this chromosome is somehow differently regulated in neural stem cells. We next focus this study on the analysis of *2xTub-GAL80^X-5B8,19E7^* line to understand why these *GAL80* insertions do not allow for complete GAL4 repression even if present at higher doses.

### *Tub-GAL80x2^X-5B8,19E7^* is not expressed in green *Drosophila* NBs

The results described above suggest a specific-brain failure of GAL80 to inhibit GAL4. To obtain more information about *GAL80* expression, we designed FISH probes that recognised *GAL80* mRNAs. The probes were tagged with a red fluophore (Methods). Analysis of *2xTub-GAL80^X-5B8,19E7^* larval brains revealed that GFP^+^ NBs lack *GAL80* RNAs FISH signals, which was not the case for GFP− NBs and surrounding cells that clearly presented red fluorescent dots (Figure 6A). These results suggest that *2xTub-GAL80^X-5B8,19E7^* expression is indeed abolished in GFP^+^ NBs. The divisions of the NBs are highly stereotyped and daughter GMCs are always generated at the same location. Thus, as the NB continues to cycle, the older daughter GMCs become more distant and placed away from the NB (Homem and Knoblich, 2012). We noticed that in some NB/GMCs clusters, localised within the central brain, the GFP-NLS signal became weaker in the GMCs placed further away from the NB. Interestingly, GMCs displaying weak GFP fluorescence were positive for *GAL80* RNAs FISH signals (Figure 6A - above the red dotted line). This observation suggests that *Tub-GAL80* expression might have been re-activated in the oldest GMCs.

**Figure 6:**
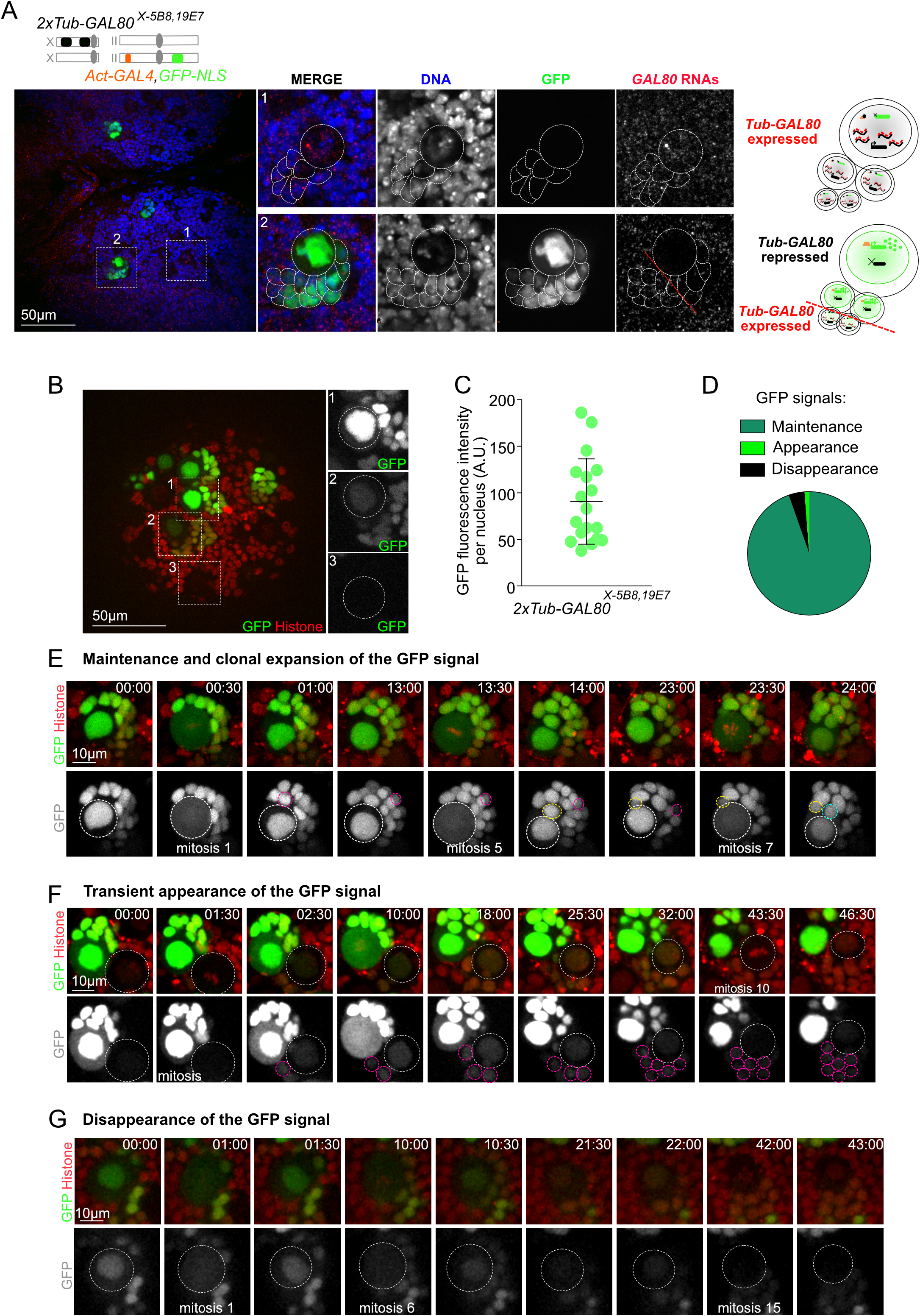
*Tub-GAL80^X-5B8,19E7^* expression is dynamic and reversible. (A) Images of whole mount brain lobes from RNA FISH experiment with probes against the GAL80 RNAs (red and grey in zoom insets) and labeled with GFP booster (green and grey in zoom insets) and DAPI for DNA (blue and grey in zoom insets). Schematic representation of cells is shown next to the images. White dotted lines surround NBs and GMCs clusters. (B) Single still from a time-lapse movie of a brain lobe (large inset) and NBs (zoom insets) expressing *Tub-GAL80^X-5B8,19E7^, Act-GAL4,GFP-NLS* (green) and *histone-RFP* (red). Several NBs from the same brain lobe present different levels of GFP intensity. (C) Dot plot showing the GFP intensity (A.U.) of NBs at the beginning of movies (n=18 NBs from 3 brains). (D) Pie chart of the different GFP dynamics (n=168 NBs from 17 brain lobes). (E-G) Stills of time-lapse movies of mitotic NBs expressing *Tub-GAL80^X-5B8,19E7^, Act-GAL4,GFP-NLS* (green) and *histone-RFP* (red) to monitor GFP and chromosome dynamics. White and colored dotted circles surround NBs and daughter GMCs, respectively. GFP signal present different dynamics: (E) maintenance and clonal expansion, (F) the transient appearance and (G) the disappearance of the GFP signal.

### *Tub-GAL80x2^X-5B8,19E7^* expression is dynamic and reversible

To obtain a dynamic view of the *Tub-GAL80* expression, we combined the same system - *Act-GAL4, GFP-NLS/2xTub-GAL80^X-5B8,19E7^* - with the expression of histone H2Av variant tagged with RFP to follow chromosome behaviour. We performed time-lapse microscopy of larval brains for up to 48hrs, which represents roughly two thirds of the total proliferative window of the central nervous system during third instar larvae-72hrs. We analysed 168 NBs from 17 brain lobes and using our imaging set up we could identify mitotic entry and exit even after long periods of laser exposure at the end of the filming period. We also did not detect nuclear fragmentation, which is a sign of apoptosis. Together, we concluded that NBs were not being subjected to deleterious phototoxicity in our imaging conditions (see methods for details).

At the start of the movie several green clusters of NBs and GMCs could be easily noticed and interestingly, showed different green fluorescent intensities. We could also identify non-fluorescent NBs as in the fixed preparations, confirming on one hand the concomitant presence of the two populations (GFP^+^ and GFP− NBs), but also possibly suggesting different birth timings revealed by the intensity of the GFP signal (Figure 6B-C). As the green NBs underwent consecutive mitosis, we noticed that the daughter GMC generated at each cell division was always green. Interestingly, we also observed that in some of the “oldest” GMCs the intensity of the green fluorescence was decreased. These results are consistent with the results of mRNA *GAL80* FISH described above and suggest that *GAL80* expression might be re-established in older GMCs. They also suggest that the young GMCs might inherit GFP-NLS through mitosis as the signal diffuses through the cytoplasm. This type of behaviour, maintenance of GFP in the NB and inheritance by the daughter GMC was observed in the large majority of all NBs presenting GFP signals at the start of the movie (n=158 out of 168 NBs) (Figure 6D-E).

In NBs that were initially non-fluorescent and so presumably expressing *GAL80*, we noticed the transient rise in GFP signal, which was also transmitted upon mitosis to the GMCs (Figure 6F). This was a much less common event as only a small proportion (1,2%, n=2 out of 168 NBs) of third instar NBs behaved in this way (Figure 6D). Importantly, these observations show that at this stage of development, while some NBs have already lost *Tub-GAL80* expression and thus accumulated a high level of GFP-NLS fluorescence, other NBs switch from a *Tub-GAL80* expressing to a *Tub-GAL80* repressive status.

Finally, a third type of behaviour was also noticed. In this case, GFP-NLS fluorescence disappeared from NBs (n=8 out of 168 NBs, Figure 6G). Interestingly, the lack of signal was maintained for many hours suggesting the maintenance of the expression of *Tub-GAL80* during this period of time. Altogether, time-lapse analysis of GFP-NLS in third instar larval brains reveals that *Tub-GAL80* expression in NBs from X-double inserted line, undergoes dynamic and to lower extend reversible changes resulting in GAL4 activity and GFP expression.

### *2xTub-GAL80^X-588,19E7^* expression is influenced by environmental changes

The dynamics and specificity of *GAL80* expression in NBs suggested a possible epigenetic regulation, specific to the brain during *Drosophila* development. This appears particular important in respect to the X chromosome since the presence of two *GAL80* copies resulted in the expression of GFP-NLS at a high frequency (Figure 5D-E). Epigenetic regulation is often used during development to provide adaptability to different environmental conditions (Friedrich et al., 2019). We tested if different stresses could influence the system using the *2xTub-GAL80^Xfor chromosome loss-5B8,19E7^* as a reporter of modifications in gene expression pattern. We controlled three parameters: (1) the food composition, (2) the abundance of parent flies and (3) the temperature of incubation and we assessed GFP signal frequency in L3 brain lobes. Importantly, each parameter was tested separately, however all conditions within each experiment were performed at the same time to allow comparison.

In all of our experimental set ups described so far, fly crosses were composed of about 20-30 parents and cultured at 25°C on a protein-rich medium, which are the standards used in *Drosophila* culture. We first altered food composition and cultured flies in a protein-poor medium made of cornmeal and low yeast content. Surprisingly, the number of GFP^+^ clusters was significantly reduced in this latter condition when compared to standard rich medium (Figure 7A). Then, as a way to influence food accessibility, we varied the number of parent flies to induce different larval crowding. The addition of <5, ±30 or >100 parents did not change GFP signal frequency (Figure 7B). Finally, we switched the temperature of larval incubation varying from 18°C to 29°C. Temperature variations strongly impacted the frequency of GFP^+^ cells. Interestingly, this was in a dose-dependent manner as the frequency gradually decreased as temperature increased (Figure 7C). Although we confirmed that change in environment conditions influences the system, it is important to mention that whatever the condition (Figure 7), the GFP signal was highly variable within a define experimental set up, as we described above (Figure 2 and Supplementary Figure 3). This suggests that abundant and yet unknown parameters influence the system.

**Figure 7:**
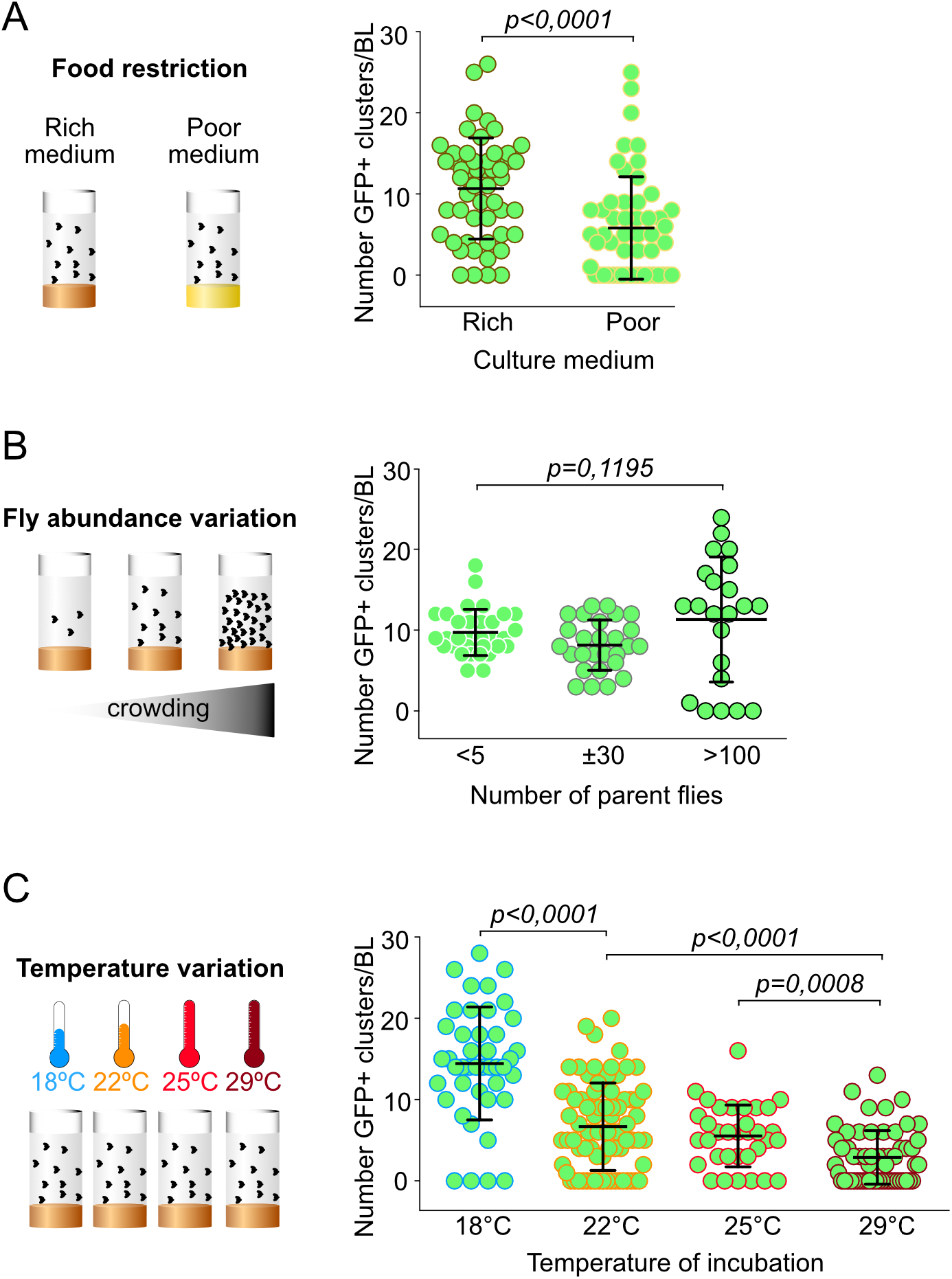
Environmental changes influence the expression/repression of *Tub-GAL80*. (A-C) Schematic representation of the different environmental stresses and dot plot showing the number of GFP^+^ clusters/brain lobe in larvae (A) raised on different culture media-proteinrich (n=70 BLs from 35 brains) or-poor medium (n=62 BLs from 31 brains), in tubes (B) containing <5 (n=32 BLs from 16 brains), ±30 (n=137BLs from 14 brains) or >100 (n=22 BLs from 11 brains) parent flies and (C) incubated at different temperatures-18ºC (n=43 BLs from 22 brains), 22ºC (n=76 BLs from 38 brains), 25ºC (n=34 BLs from 17 brains) or 29ºC (n=58 BLs from 29 brains). Statistical significance was determined by a Mann-Whitney test and p corresponds to the p-value.

Together, these results suggest that NBs of the *Drosophila* developing larval brain undergo a novel gene expression regulation mode that appears to be dynamic, reversible and capable of being influenced by different environmental conditions. It will be interesting to investigate if the findings described here also apply to other more complex central nervous systems.

## Discussion

Here we sought to develop a new probe to monitor chromosome loss. Using the bipartite UAS-GAL4 reporter system in combination with the GAL80 repressor, we reasoned that loss of the *GAL80* containing chromosome would be translated by the appearance of nuclear GFP signals. While this reasoning appears to apply to some larval tissues, including imaginal discs, it appears nonfunctional in the brain. Quite surprisingly, however this study revealed an unexpected level of regulation of gene expression in the larval brain.

The UAS-GAL4/GAL80 system is widely used among the *Drosophila* community to spatially and temporally control gene expression (McGuire et al., 2003). It was therefore difficult to predict the results obtained in the *Drosophila* developing brain and described here. Our findings on the lack of *GAL80* expression highlights a weakness of the system and precludes, or at least alerts, its use in the *Drosophila* developing brain for MARCM or “gypsy-trap” analysis that are based on the *GAL80* sequence loss after heat-induced mitotic recombination (Lee and Luo, 1999) or on the gypsy dependent *GAL80* repression (Li et al., 2013).

One of the most surprising result was related with the variability of cell types becoming GFP^+^ according to the *GAL80* insertion position on each chromosome. For instance, while in lines *Tub-GAL80^X-5B8^* or *Tub-GAL80^III-92F3^* the majority of green cells were NBs and GMCs, in line *Tub-GAL80^III-82A1^* we mainly detected green neurons or green glial cells. These results suggest that different chromosome regions within the fly genome might be subjected to particular rules of gene regulation in a cell type (brain specific)-dependent manner, as already suggested for heterochromatin-mediated silencing during larval development (Lu et al., 1996). Additionally, it is important to consider that for the same line, the position of green cells was also quite variable. For example, in line *Tub-GAL80^III-70C3^*, green NBs were positioned at different regions of the central brain, in different brain lobes. These results, combine with the results obtained by time-lapse microscopy suggest a certain randomness or stochasticity of the system.

The lack of repression capacity found for most *GAL80* insertions in different chromosome locations for the X, II and III chromosomes suggest however that one single *GAL80* copy is not sufficient to inhibit GAL4 activity. These results can be explained by an imbalance in terms of GAL4/GAL80 stoichiometry (Pfeiffer et al., 2010). Generally, the strength of gene expression relies on the strength of its promoter. The stoichiometry imbalance can thus be explained by the different promoters used in this study. Indeed, the lines used here: *Tub-GAL80* and *Act-GAL4*, even if considered as ubiquitous and containing strong promoters seem to be rather different. The differences in promoter strength seem to be supported by our findings showing that when both *GAL4* and *GAL80* sequences were under the same actin promoter, the frequency of GFP^+^ cells was drastically reduced in the brain. However, even if plausible, this explanation might also be quite simplistic and does not explain all our results. While lines *Tub-GAL80x2^II-22A,53B2^* and *Tub-GAL80x2^III-62E1,96F3^*, which contained two copies of *Tub-GAL80* did repress GAL4 activity and represent good candidates as probes for chromosome loss, lines *Tub-GAL80x2^X-5B9,19E7^, Tub-GAL80x2^II-25A3,59D3^* and *Tub-GAL80x2^III-65B2,99F^* did not suppress all GFP^+^ cells. This was even more obvious in line *Tub-GAL80x2^X-5B9,19E7^*, on the X chromosome, which suggests that the chromosome X might be subjected to an even more particular and yet unknown gene regulation process.

From all the lines analysed in this study, the X chromosome appears as a particular chromosome in terms of gene regulation specifically in NBs. The presence of two or four *Tub-GAL80* copies did not repress GFP expression. GFP was inherited by the daughter GMCs during mitosis, suggesting a dynamic behavior. Importantly, older GMCs from a NB/GMCs cluster were frequently less green than younger GMCs or even the mother NB. Together with the RNA FISH analysis showing *GAL80* mRNAs signals in GMCs away from the NB, these results suggest that *GAL80* expression might be re-installed as GMCs age and become more differentiated. These observations indicate therefore that the particular and most likely random gene regulation of the X chromosome is associated with neural stem cell identity.

The findings that different *Tub-GAL80* lines generate different GFP^+^ patterns exclusive in the brain suggest a specific regulation at the chromatin level most likely dependent on epigenetic marks that might allow adaptation in adverse situations. The plasticity of the system seems to be partially supported by our observations of different green cell frequency in response to different environmental changes. For example, the difference in the frequency of GFP^+^ NBs observed at lower temperature might also reflect a safeguard mechanism that closes chromatin (less GAL80) to slow down neurogenesis. In agreement, it has been proposed that neurogenesis requires a certain neuroblast plasticity and competence to generate different types of neurons (Pearson and Doe, 2003; Cleary and Doe, 2006). This plasticity at the scale of an organ like the brain can positively serve adaptation and evolution. In light of this possibility, the epigenetic plasticity, inherent and specific to the brain might allow a rapid adaptation to stress and environmental changes.

## Acknowledgments

We acknowledge the PiCT-IBiSA platform and Nikon Imaging Center at Institut Curie for image set up. We thank V. Marthiens, S. Gemble, M. Bud zyk, F. Leulier, J. Merlet, P. Tran and the Bardin team for helpful discussion and/or comments on the manuscript. This work was supported by an ERC CoG (ChromoNumber - LS3, ERC-2016-COG), Institut Curie and the CNRS. A.G was funded by FRM (ECO20170637529) and LLCC (IP/SC-16533) fellowships.

## Author contribution

A.G performed all experiments, analyzed the data, generated the figures and wrote the paper. C.P generated certain tools and A.S helped for some experiments. C.P and A.S helped with fly pushing. P.S generated the *Tub-GAL80* plasmid. R.B conceived and supervised the project. A.G, R.B and A.B interpreted the data which was discussed between all authors during the preparation of the manuscript.

## Materials and Methods

### Fly husbandry and fly stocks

For most experiments, flies were raised in plastic vials containing homemade standard *Drosophila* rich culture medium (0.75% agar, 3.5% organic wheat flour, 5% yeast, 5.5% sugar, 2.5% nipagin, 1% penicillin-streptomycin (Gibco #15140), and 0.4% propanic acid). Fly stocks were maintained at 22ºC and experimental crosses at 25ºC. For food restriction experiment, flies were raised on homemade protein-poor medium (0.75% agar, 7% cornmeal, 1.4% yeast, 5.2% sugar, 1.4% nipagin) at 22ºC and compared to flies raised on homemade standard rich medium at 22ºC. For temperature variation experiment, flies were laying eggs for 24hours and tubes containing progeny were maintained at 18ºC, 22ºC, 25ºC or 29ºC for 7, 5, 5 or 4 days, prior dissection, respectively. For the MARCM experiment, fly crosses were kept at 22ºC. L2 progenies were heat-shocked 1hour at 37 ºC in a water bath and maintained at 22ºC for 30hours before dissection.

Fly stocks are summarized in Supplementary Table 1.

### *GAL80 Drosophila* lines establishment

The plasmid containing codon-optimized *GAL80* sequence driven by a tubulin promoter is a gift from Allison Bardin and corresponds to the combination of pattB-tubP-SV40 - generated by Lee and Luo (Lee and Luo, 1999)-with the codon optimized GAL80 sequence from pBPGAL80Uw-6-a gift from Gerald Rubin (Addgene plasmid #26236, http://n2t.net/addgene:26236; RRID:Addgene-26236) (Pfeiffer et al., 2010). Then, the plasmid was inserted in a P[acman] vector and send to Bestgene^®^ company to integrate it into the *Drosophila* genome at specific insertion sites using PhiC31 integrase-mediated transgenesis system.

To obtain *Drosophila* recombinants carrying two copies of *Tub-GAL80* on the same chromosome, we used female meiotic recombination and selected recombination events based on the fly eyes colour, a method widely used to generate *Drosophila* recombinants. Indeed, all *Tub-GAL80* are associated with the white^+^ transgene expressing marker, which is used as a marker for efficient transgene insertion as it confers yellow to red eye colour. Simply, in the presence of two copies, as for efficient recombination, fly eyes display a strong red colour.

### Immunofluorescence of *Drosophila* larval whole mount tissues

Mid third-instar larval (L3) brains and imaginal discs were dissected in fresh Phosphate-Buffered Saline 10X (PBS, VWR #L182-10) and fixed for 30 minutes (min) at room temperature (RT) in 4% paraformaldehyde (EMS # 15710) diluted in PBS. Fixed tissues were washed and permeabilized three times 15 min in PBST3 or PBST1 (PBS, 0,3% or 0,1% Triton X-100, Euromedex #2000-C). For antibody staining, larval tissues were incubated in primary antibodies diluted in PBST3 or PBST1 overnight at 4ºC in a humid chamber. After 3×15 min washes in PBST3 or PBST1, tissues were incubated in secondary antibodies diluted in PBST3 or PBST1, O/N at 4ºC and protected from light in a humid chamber. Tissues were then washed 3×15 min in PBST3 or PBST1, rinsed in PBS and mounted between slides (Thermo Fisher Scientific #AA00008232E00MNT10) and 12-mm circular cover glasses (Marienfield Superior #0111520) with 5μl of homemade mounting medium (1,25% n-propyl gallate, 75% glycerol, 25% H_2_O).

For GFP screening of larval brains and imaginal discs, one step of O/N incubation at 4ºC with GFP booster (1:250, Alexa Fluor ^®^ 488 Chromotek #gb2AF488), Phalloidin-647 (1:250, Thermo Fisher Scientific #A-22287) and DAPI (1:1000, Thermo Fisher Scientific #62248) were performed followed by 3×15 min washes in PBST3, a rinse in PBS and mounting.

Primary antibodies used in this study are: chicken (chk) anti-GFP (1:1000, Abcam #ab13970), guinea pig (GP) anti-Deadpan (Dpn) (1:1000, J. Skeath), mouse anti-Prospero (1:500, MR1A, DSHB), rat anti-Elav (1:100, 7EA10, DSHB), mouse anti-Repo (1:500, 8D15, DSHB), rabbit (Rb) anti-Sas4 (1:500, Basto et al 2006), GP anti-Centrosomin (Cnn) (1:1000, E. Lucas and J.W.R.). Secondary antibodies (1:250) used in this study are: chk-488 (Thermo Fisher Scientific #A-11039), Rat-546 (Thermo Fisher Scientific #A-11081), Rb-568 (Thermo Fisher Scientific #A-10042), mouse-546 (Thermo Fisher Scientific #A-11030) and GP-647 (Thermo Fisher Scientific #A-21450).

Images were acquired with 40x (NA 1.25), 63x (NA 1.32) or 100x (NA 1.4) oil objectives on a wide-field Inverted Spinning Disk Confocal Gattaca/Nikon (a Yokagawa CSU-W1 spinning head mounted on a Nikon Ti-E inverted microscope equipped with a camera complementary metal-oxide semiconductor 1.200 x 1.200 Prime95B; Photometrics). Intervals for z-stack acquisitions were set to 0.5 to 1.5μm using Metamorph software.

### Live imaging of *Drosophila* larval brains

Mid second-instar larval (L2) brains were dissected in Schneider’s *Drosophila* medium (Gibco #21720024) supplemented with 10% heat-inactivated fetal bovine serum (Gibco #10500), penicillin (100 U/ml) and streptomycin (100 μg/ml). Several brains were placed in 10μl of medium on a glass-bottom dish (Dutcher #627870), covered with a permeable membrane (Standard YSI), and sealed around the membrane borders with oil 10 S Voltalef (VWR Chemicals). Images were acquired with 60x oil objective (NA 1.4) on two microscopes: an Inverted Spinning Disk Confocal Roper/Nikon (a Yokagawa CSU-X1 spinning head mounted on a Nikon Ti-E inverted microscope equipped with a camera EMCCD 512 x 512 Evolve; Photometrics) and the wide-field Inverted Spinning Disk Confocal Gattaca/Nikon (a Yokagawa CSU-W1 spinning head mounted on a Nikon Ti-E inverted microscope equipped with a camera complementary metal-oxide semiconductor 1.200 x 1.200 Prime95B; Photometrics), controlled by Metamorph software. For both microscopes, images were acquired at time intervals spanning 30 min and 50 z-stacks of 1.5 μm.

### DNA Fluorescent *in situ* hybridization

After fixation, permeabilization and O/N incubation with GFP booster (description below), brains were washed 3×15min in PBT3 and fixed a second time 30 min in 4%PFA. Then, brains were rinse 3x in PBS, washed 1×5 min in 2xSSCT (2X Saline Sodium Citrate (Euromedex #EU0300-A) 0,1% Tween-20 (Sigma Aldrich #P1379) diluted in water), 1×5 min in 2xSSCT/50% formamide (Sigma Aldrich #47671), transferred in pre-warmed 2xSSCT/50% formamide and pre-hybridized 3 min at 92°C. In the meantime, DNA probes diluted in the Hybridization Buffer (20%dextransulfate (Sigma Aldrich #D8906), 2xSSCT, 50%formamide, 0,5mg/ml salmon DNA sperm (Sigma Aldrich #D1626)) were denature at 92°C. After removal of the supernatant, brains were incubated in the probes solutions and hybridize 5 min at 92°C and O/N at 37°C. Brains were then rinse at RT, washed 1×10 min at 60°C and 1×5 min at RT in 2xSSCT. Finally, after a rinse in PBS brains were mounted as described below.

DNA probes used in this study were against chromosomes X (80ng/μl), II (40ng/μl) and III (80ng/μl).

### RNA Fluorescent *in situ* hybridization

L3 brains were dissected in fresh PBS, fixed 30 min in 4% formaldehyde (EMS # 15686) and washed and permeabilized in PBSTw (PBS 0,3% Tween-20). Brains were incubated with GFP booster and Phalloidin diluted in PBSTw, O/N at 4°C in a humid chamber. After 3×15 min washes PBSTw, RNA hybridization was performed as described by Yang and colleagues (Yang et al., 2017). RNA probes against *GAL80* were designed by the biosearchtech^®^ technical support team (https://www.biosearchtech.com/) and labeled with quasar 570.

### Polymerase Chain Reaction (PCR) of *Drosophila* lines

Good quality genomic DNA was extracted according to the protocol provided by the Vienna Drosophila Resource Center from flies heterozygous for *Tub-GAL80^II-53B2^* (one *GAL80* copy), homozygous for *Tub-GAL80^II-53B2^* (two *GAL80* copies) and heterozygous recombinant *2xTub-GAL80^X5B8,19E7^* (two *GAL80* copies). PCR analysis was performed using the following primers: *GAL80* forward (5’ CGGTGCCGAATGCTGCTCCCA 3’) and *GAL80* reverse (5’ CCGAACGTGGTGGTCACCAGA 3’).

---

**Supplementary Figure 1:**
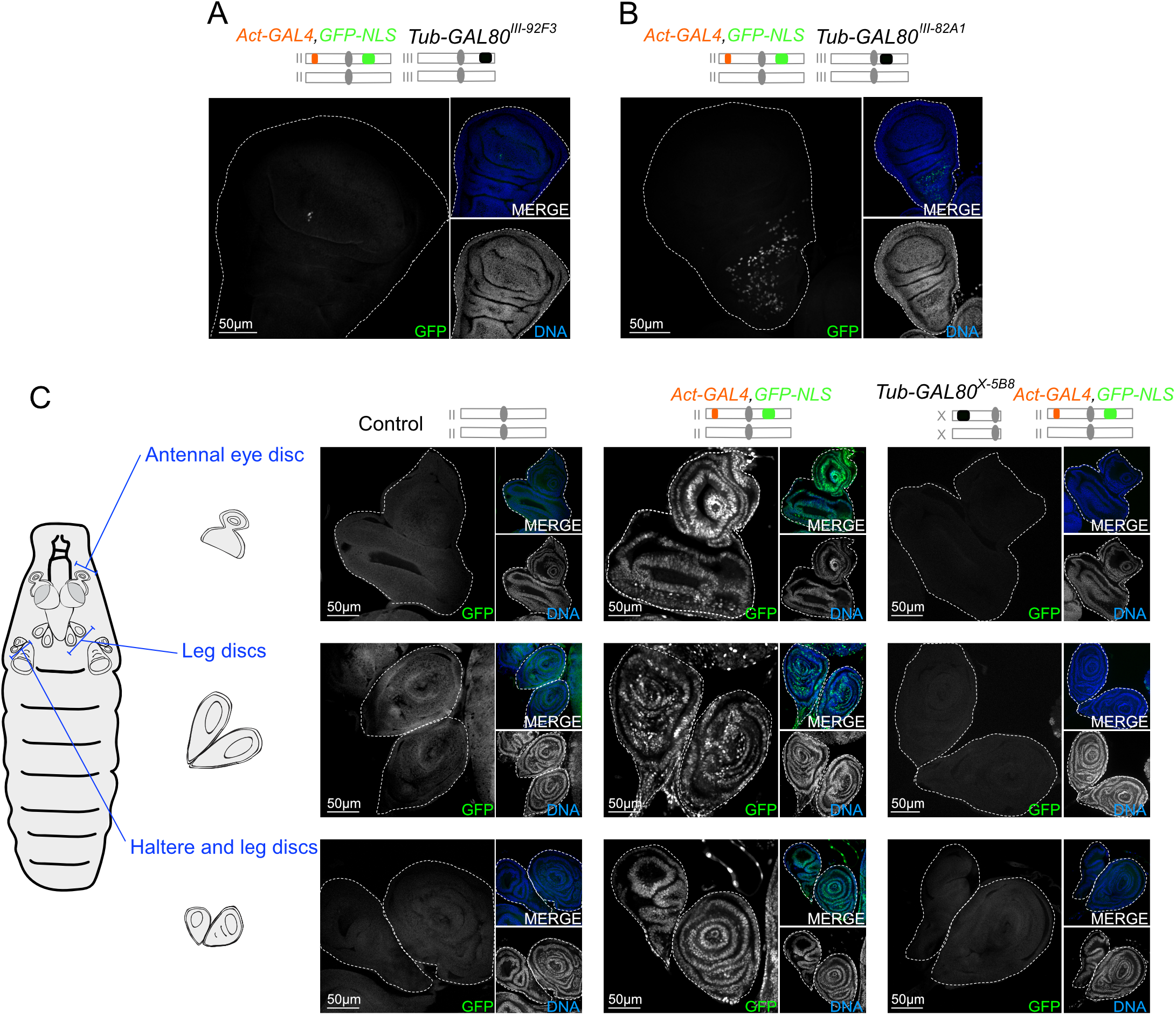
The GAL4/GAL80 system is functional in imaginal discs of the *Drosophila* larvae. (A-C) Images of whole mount imaginal discs labeled with GFP booster (grey and green in large and small insets, respectively) and DAPI for DNA (blue and grey in small insets). White dotted lines surround discs. Schematic representation of the genotypes is shown above the images. Few wing discs presented low (A) or high (B) levels of GFP^+^ cells. (C) Schematic representation of imaginal discs in *Drosophila* larvae. Control discs present no GFP signal. In the presence of the activator GAL4, *GFP-NLS* is expressed and all cells present GFP^+^ nuclei. GAL80 efficiently inhibits GAL4 and thus, cells are GFP negative.

**Supplementary Figure 2:**
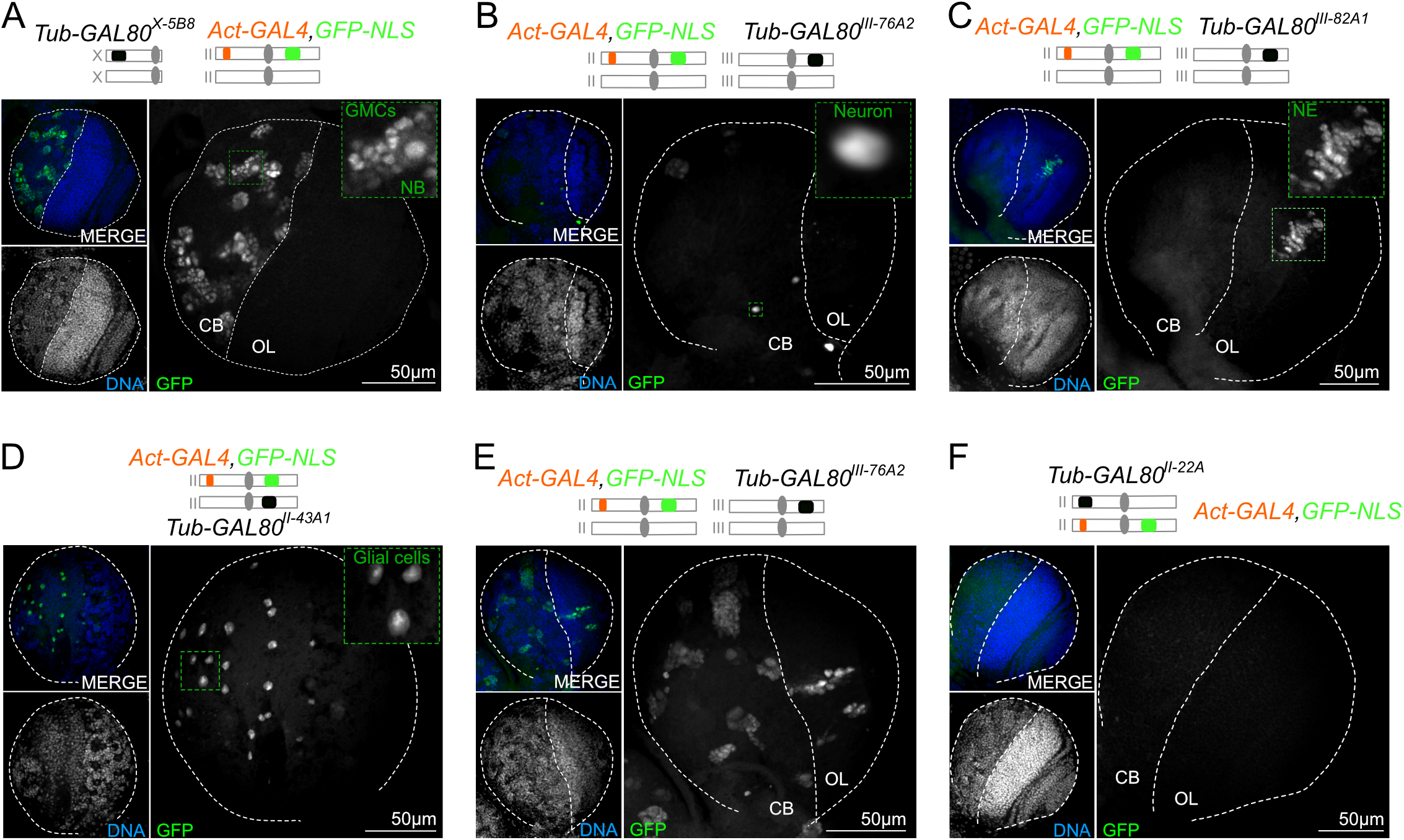
The GFP-NLS signal is sufficient to distinguish all cell types. (A-F) Images of whole mount brain lobes labeled with GFP booster (grey and green in large and small insets, respectively) and DAPI for DNA (grey and blue in small insets). Schematic representation of the genotypes is shown above the images. White dotted lines delimitate brain lobes and separate CB and OL. GFP^+^ cells are (A) NBs with GMCs, (B) individual neurons, (C) NE cells, (D) glial cells, or (E) a mix population of different cell types. (F) Very few brain lobes presented an absence of GFP signal.

**Supplementary Figure 3:**
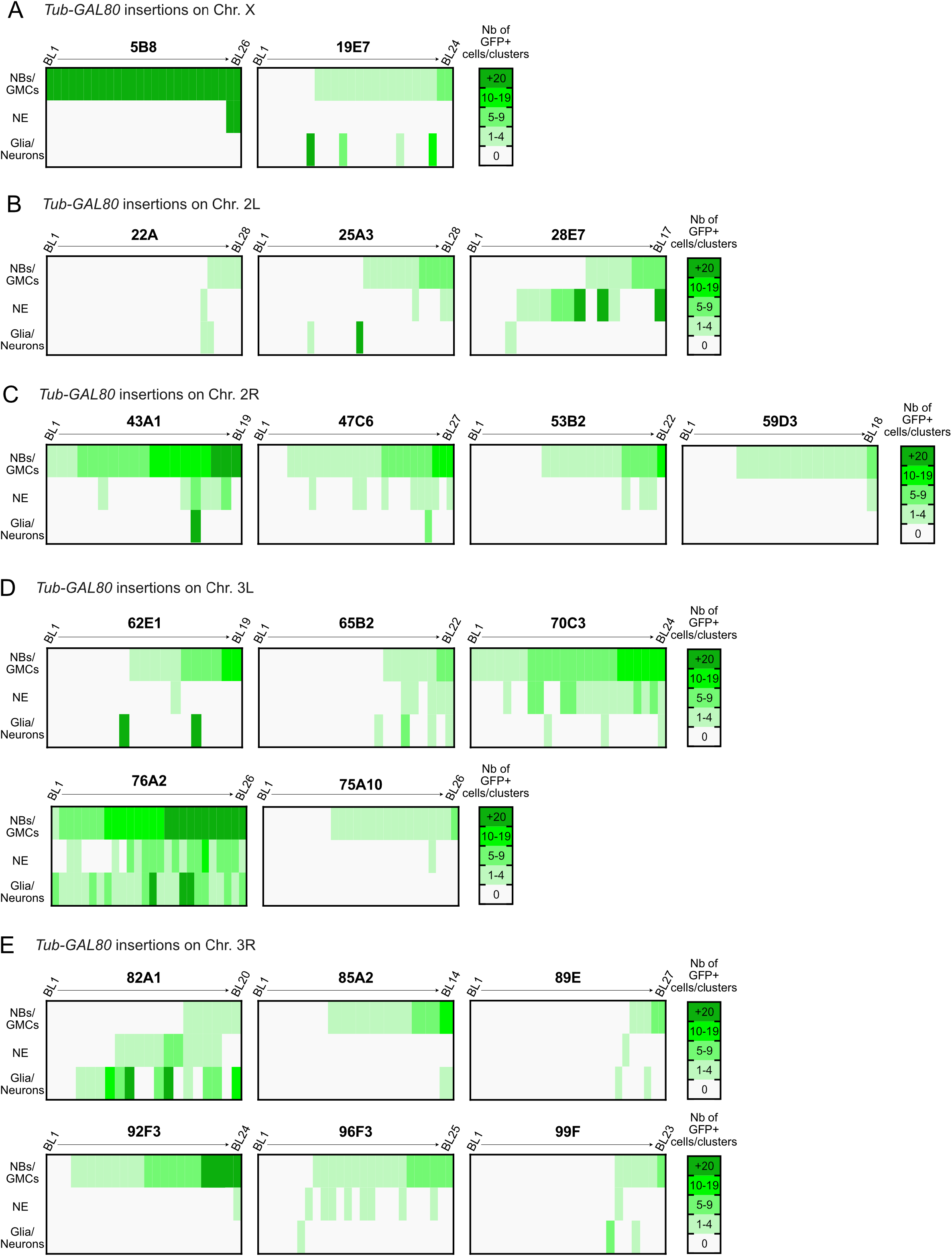
The majority of the brains present GFP signals which show variable frequency. (A-E) Heat map showing the number of GFP^+^ (green color code) cells for glial cells and individual neurons and GFP^+^ clusters for NBs with GMCs and NE cells. All brain lobes analyzed are numerated (BL_1_ → BL_n_) and the *Tub-GAL80* insertion sites are indicated above the graphs. The cell type and the number of GFP^+^ cells/clusters are highly variable between all conditions with *Tub-GAL80* inserted on the (A) X chromosome, on the (B) left and (C) right arms of the chromosome II and on the (D) left and (E) right arms of the chromosome III (n=14 to 28 BLs/ condition). Representative images are shown in Figure 2 and Supplementary Figure 2.

**Supplementary Figure 4:**
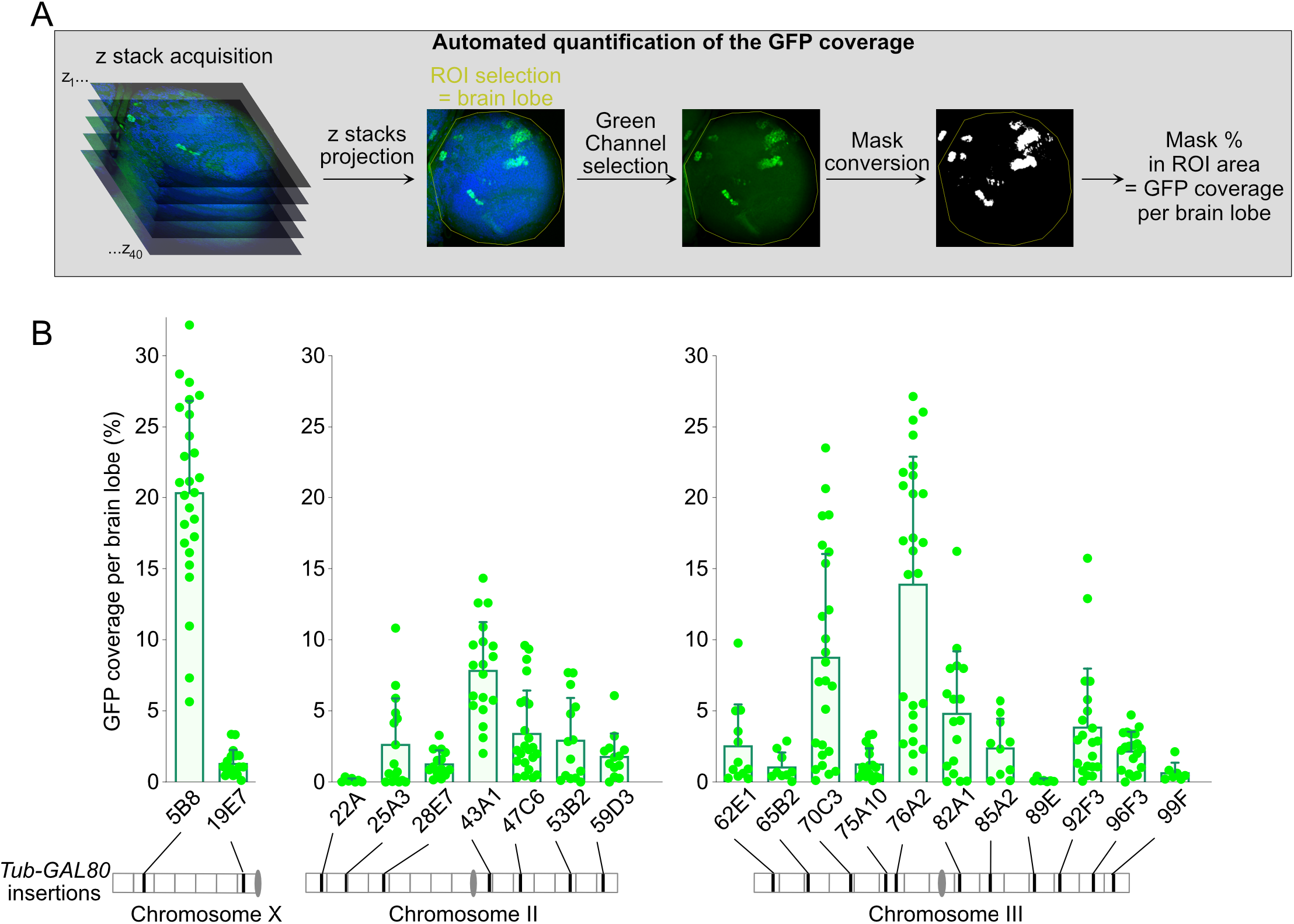
The coverage of the GFP signals is variable between all GAL80 conditions. (A) Schematic representation of the automated quantification of the GFP coverage. ROI corresponds to “Region Of Interest”. (B) Scatter plot with bars showing the GFP coverage per brain lobe for each GAL80 condition (n=7 to 26 GFP^+^ brain lobes/condition).

**Supplementary Figure 5:**
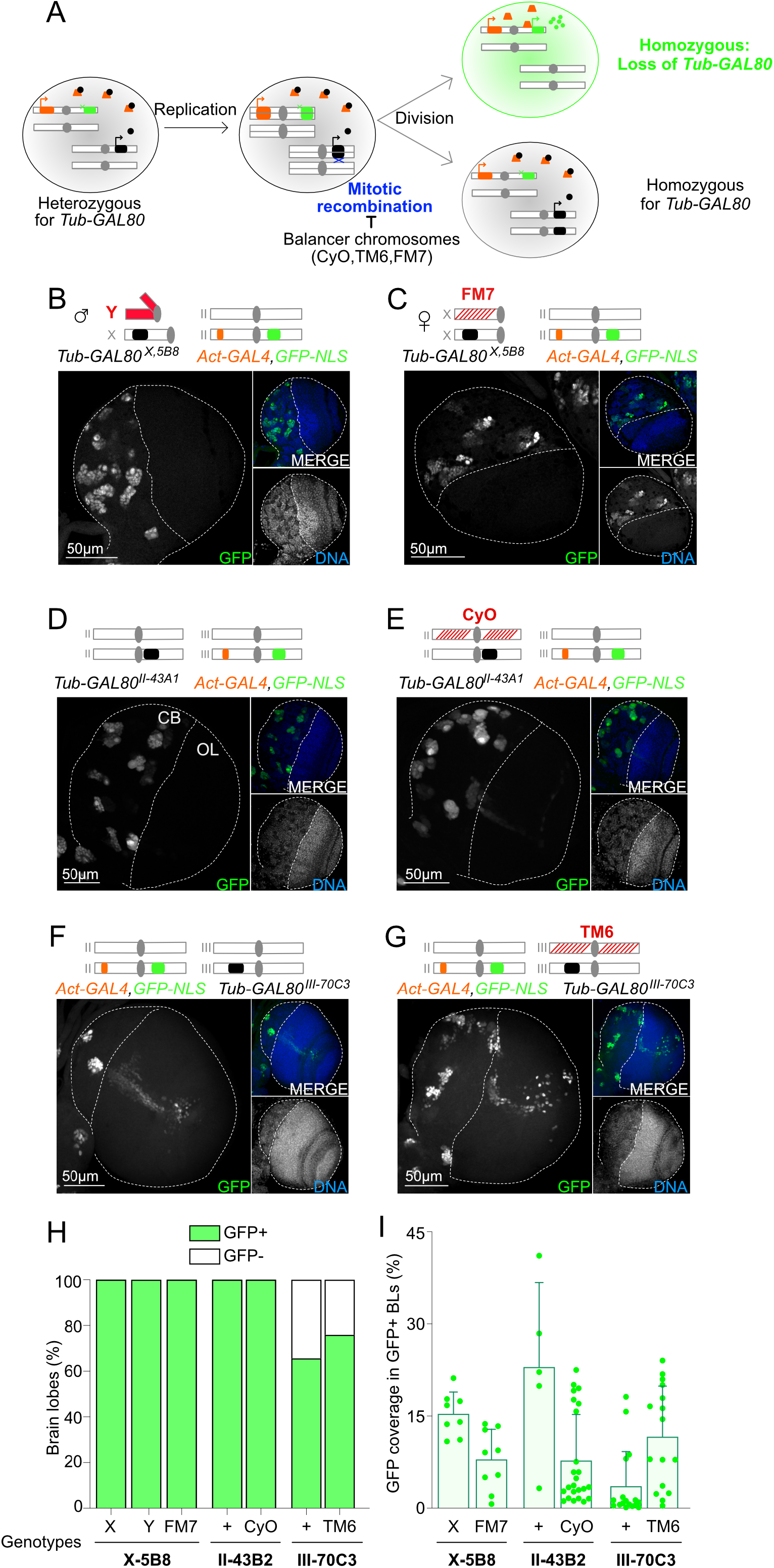
Mitotic recombination does not account for GFP appearance. (A) Schematic representation of mitotic recombination inducing loss of *Tub-GAL80* heterozygosity and GFP appearance. (B-G) Images of whole mount brain lobes labeled with GFP booster (grey and green in large and small insets, respectively) and DAPI for DNA (blue and grey in small insets). White dotted lines surround brain lobes and delimitate CB and OL. Schematic representation of the genotypes is shown above the images. (H) Graph bar showing the percentage of brain lobes without (white) and with GFP signal (green) (n=6 to 32 brain lobes/condition). (I) Scatter plot with bars showing the GFP coverage per brain lobe for each condition (n=5 to 33 GFP^+^ brain lobes/condition).

**Supplementary Figure 6:**
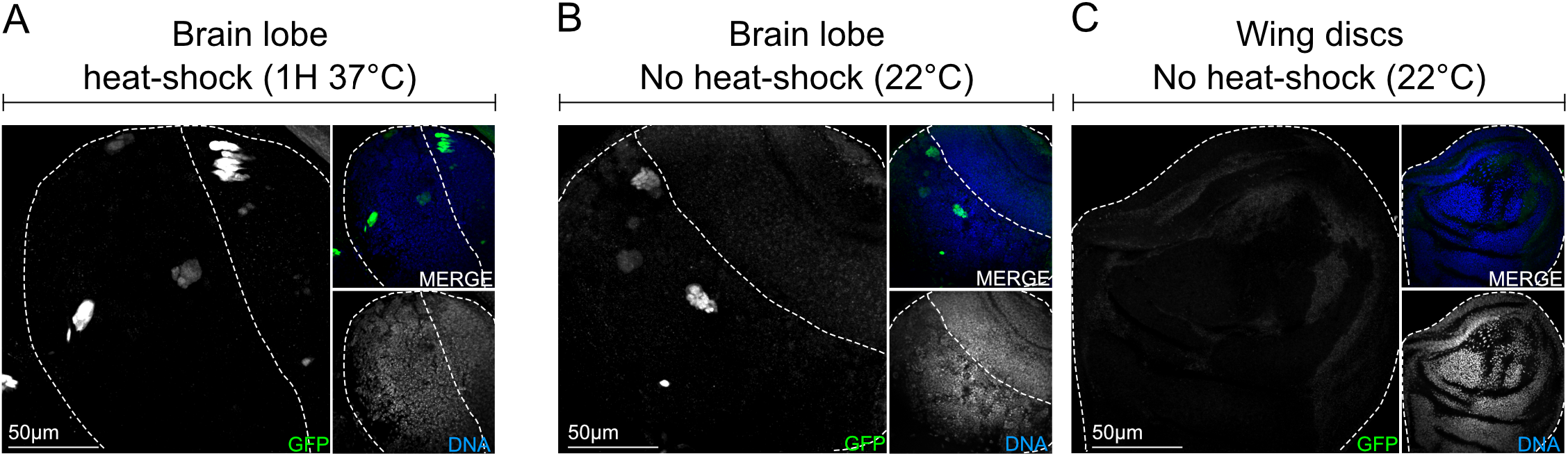
GFP^+^ NB/GMCs clones are formed in MARCM flies in absence of heat shock. (A-C) Images of whole mount *hs-FLP/+, UAS-GFP-NLS/+; Tub-GAL4, FRT82B, Tub-GAL80/FRT82B, sas4^mut^* brain lobes (A-B) or wing disc (C) labeled with antibodies against GFP (grey and green in large and small insets, respectively) and DAPI for DNA (grey and blue in small insets). White dotted lines delimitate tissues. (A) Presence of GFP^+^ clones in brain lobes of flies heat-shocked at 37ºC for 1 hour. (B-C) In the absence of heat-shock, (B) brain lobes present GFP^+^ clones, in contrast to (C) wing discs that are GFP−.

**Supplementary Figure 7:**
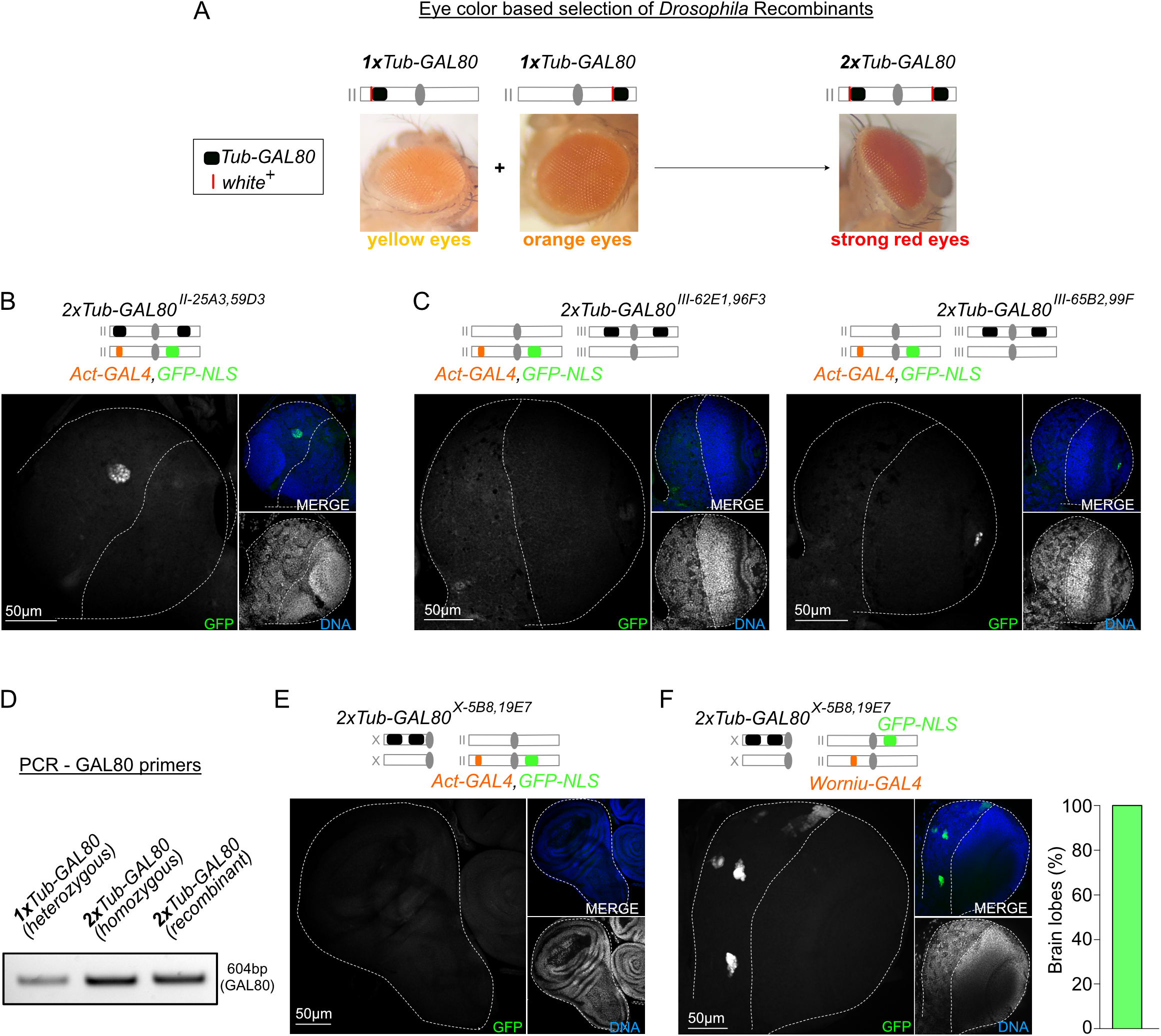
Double insertions of *Tub-GAL80* decrease GFP frequency. (A) Pictures of *Drosophila* recombinant selection to obtain lines carrying two copies of the *Tub-GAL80* cassette (black boxes) based on eyes color (white^+^ transgene - red lines). (B-C) Brain lobes labeled with GFP booster (grey and green in large and small inserts, respectively) and DAPI for DNA (blue and grey in small insets). White dotted lines surround brain lobes. Schematic representation of the genotypes is shown above the images. (D) PCR analysis of *Drosophila* lines using primers recognizing the two extremities of the *GAL80* sequence (604 base pairs). The recombinant flies (*2xTub-GAL80^X-5B8,19E7^*) present a similar or stronger band than flies carrying a single *Tub-GAL80* copy in homozygous (*2xTub-GAL80^II-53B2^*) or heterozygous (*1xTub-GAL80^II-53B2^*) state, respectively. (E) Wing disc and (F) brain lobe labeled with GFP booster (grey and green in large and small inserts, respectively) and DAPI for DNA (blue and grey in small insets) and graph bar showing the percentage of brain lobes presenting GFP^+^ cells in flies expressing *2xTub-GAL80^X-5B8,19E7^; worniu-GAL4/UAS-GFP-NLS* (n=30 brain lobes).

**Table 1 -.** List of *Drosophila* stocks

| Stock Name | Genotype | Chromosomal location | Origins |
| --- | --- | --- | --- |
| WT | w[118] |  | RRID:BDSC_5905 |
| Tub-GAL80 | alphaTub84B-GAL80 | X-5B8 | This study |
| Tub-GAL80 | alphaTub84B-GAL80 | X-19E7 | This study |
| Tub-GAL80 | alphaTub84B-GAL80 | 2L-22A3 | Bardin lab |
| Tub-GAL80 | alphaTub84B-GAL80 | 2L-25A3 | This study |
| Tub-GAL80 | alphaTub84B-GAL80 | 2L-28E7 | Bardin lab |
| Tub-GAL80 | alphaTub84B-GAL80 | 2R-43A1 | This study |
| Tub-GAL80 | alphaTub84B-GAL80 | 2R-47C6 | This study |
| Tub-GAL80 | alphaTub84B-GAL80 | 2R-53B2 | This study |
| Tub-GAL80 | alphaTub84B-GAL80 | 2R-59D3 | This study |
| Tub-GAL80 | alphaTub84B-GAL80 | 3L-62E1 | This study |
| Tub-GAL80 | alphaTub84B-GAL80 | 3L-65B2 | This study |
| Tub-GAL80 | alphaTub84B-GAL80 | 3L-70C4 | This study |
| Tub-GAL80 | alphaTub84B-GAL80 | 3L-75A10 | This study |
| Tub-GAL80 | alphaTub84B-GAL80 | 3L-76A2 | This study |
| Tub-GAL80 | alphaTub84B-GAL80 | 3R-82A1 | This study |
| Tub-GAL80 | alphaTub84B-GAL80 | 3R-85A2 | This study |
| Tub-GAL80 | alphaTub84B-GAL80 | 3R-89E11 | Bardin lab |
| Tub-GAL80 | alphaTub84B-GAL80 | 3R-92F1 | This study |
| Tub-GAL80 | alphaTub84B-GAL80 | 3R-96F3 | This study |
| Tub-GAL80 | alphaTub84B-GAL80 | 3R-99F8 | Bardin lab |
| Act-GAL4 | Actin5C-GAL4 | 2 | RRID:BDSC_4414 |
| Act-GAL4 | Actin5C-GAL4 | 3 | RRID:BDSC_3954 |
| GFP-NLS | UAS-GFP-NLS | 3 | RRID:BDSC_4776 |
| GFP-NLS | UAS-stinger | 2 | RRID:BDSC_84277 |
| Tub-GAL80 | hsFLP,alphaTub84B-GAL80,FRT19A | X-1C2 | RRID:BDSC_5132 |
| Tub-GAL80 | alphaTub84B-GAL80,FRT80B | 3L-75E1 | RRID:BDSC_5191 |
| Tub-GAL80 | alphaTub84B-GAL80 | 3 | RRID:BDSC_9490 |
| Act-GAL80 | Actin5C(loxp.GAL80.stop)loxA::p65 | 2-25C6 | RRID:BDSC_67092 |
| Tub-GAL4,FRT82B,Tub-GAL80 | alphaTub84B-GAL4,FRT82B,alphaTub84B-GAL80 | 3-79A2,82B2 | RRID:BDSC_30036 |
| hsFLP | Hsp70-FLP | X | N/A |
| FRT82B,sas4mut | FRT82B, sas4mut | 3 | Raff lab |
| Histone-RFP | HistoneH2Av-mRFP | 2 | RRID:BDSC_23651 |
| Act-GAL4,GFP-NLS/CyoGFP |  | 2 | This study |
| Act-GAL4,GFP-NLS/TM6,Tb |  | 3 | This study |
| Act-GAL4,GFP-NLS,Histone-RFP/CyoGFP |  | 2 | This study |
| GFP-NLS;Tub-GAL4,FRT82B,Tub-GAL80 |  | 2;3 | This study |
| hsFLP/FM7,kr;;FRT82B,sas4mut/TM6,Tb |  | X;;3 | This study |
| 2xTub-GAL80 |  | X-5B8,19E7 | This study |
| 2xTub-GAL80 |  | 2-22A,53B2 | This study |
| 2xTub-GAL80 |  | 2-25A3,59D3 | This study |
| 2xTub-GAL80 |  | 3-62E1,96F3 | This study |
| 2xTub-GAL80 |  | 3-65B2,99F8 | This study |
| 2xTub-GAL80;;Act-GAL4,GFP-NLS |  | X-5B8,19E7;;3 | This study |

